# Preclinical evaluation of antisense oligonucleotide therapy in a mouse model of *HNRNPH2*-related neurodevelopmental disorder

**DOI:** 10.1101/2025.11.04.686541

**Authors:** Ane Korff, Xiaojing Yang, Ozan Ozdemir, Ananya Samanta, Yong-Dong Wang, Tushar Patni, Alfonso J. Lavado, Anoop Murthy Kavirayani, Joseph Ochaba, Berit Powers, C. Frank Bennett, Hong Joo Kim, J. Paul Taylor

**Author notes:** **Corresponding author:** J. Paul Taylor, 262 Danny Thomas Place, Memphis, TN, USA 38105.

## Abstract

Mutations in *HNRNPH2* cause an X-linked disorder characterized by developmental delay, intellectual disability, motor and gait disturbances, and seizures. Murine models that reproduce key clinical features of HNRNPH2-related neurodevelopmental disorder suggest that it may result from a toxic gain of function of the mutant protein or a complex loss of normal HNRNPH2 function with impaired compensation by its homolog, HNRNPH1. In this study, we tested gapmer antisense oligonucleotides (ASOs) that target murine *Hnrnph2* in a non-allele-specific manner. The lead ASO reduced *Hnrnph2* mRNA and protein levels while inducing compensatory upregulation of *Hnrnph1* in both WT and *Hnrnph2* mutant mouse brains. A single intracerebroventricular injection of the *Hnrnph2* ASO into neonatal mutant *Hnrnph2* mice rescued molecular and audiogenic seizure phenotypes and improved motor and cognitive functions. ASO treatment at the juvenile stage also rescued audiogenic seizures and motor deficits. In contrast, *Hnrnph2* ASO administration did not improve survival, body weight, or hydrocephalus. In human iPSC-derived neurons, a human-specific *HNRNPH2* research ASO reduced *HNRNPH2* and upregulated *HNRNPH1* mRNA levels. Mechanistically, we demonstrate that *HNRNPH1* expression is regulated by alternative splicing and that HNRNPH2 modulates this process. These findings provide preclinical proof of concept for *HNRNPH2* ASO therapy and offer insights into its underlying molecular mechanism.

**One Sentence Summary:** ASO-mediated *Hnrnph2* knockdown induces *Hnrnph1* upregulation and rescues phenotypes in a mouse model of HNRNPH2-related neurodevelopmental disorder.

## Introduction

*HNRNPH2*-related disorder is a rare neurodevelopmental disorder (NDD) caused by mutations in *HNRNPH2*, which is located on the X chromosome (*1*). HNRNPH2 is part of the HNRNP F/H family of RNA-binding proteins, which also includes HNRNPH1, HNRNPH3, HNRNPF, and GRSF1. Together, these proteins play a critical role in RNA maturation by regulating alternative splicing, 5′ capping and 3′ polyadenylation of RNAs, and RNA export (*2*). HNRNP F/H proteins contain a proline-tyrosine nuclear localization sequence (PY-NLS) within their glycine-tyrosine-arginine-rich (GYR) domain, which binds karyopherin β2 (Kapβ2) to regulate their nucleocytoplasmic transport (*3*).

The first six *HNRNPH2*-NDD cases reported in 2016 were all females carrying one of three de novo mutations. Subsequent studies have expanded the genotypic spectrum to include 11 distinct de novo variants (*4*) as well as some maternally inherited cases (*5, 6*). Furthermore, although *HNRNPH2* mutations were initially thought to be embryonically lethal in males, several male patients have since been identified (*7–9*), but they remain in the minority. Most individuals identified to date carry a nonsynonymous single nucleotide variant within or adjacent to the NLS of HNRNPH2, with the 2 most common missense variants, R206W and R206Q, located within the NLS. Another variant located within the NLS, P209L, has been found in only one patient to date.

The phenotypic spectrum of *HNRNPH2*-NDD varies considerably between individuals. This can be attributed in part to sex/dosage effects and the location of specific mutations (*4, 6, 10*). However, other factors may play a role, as female patients carrying the same variant show considerable differences in their symptom severity. In this regard, skewed X-inactivation has emerged as a possible contributor to phenotypic variation (*5*). Most *HNRNPH2*-NDD patients exhibit some combination of developmental delay, intellectual disability, language impairment, motor function deficits, and growth and musculoskeletal problems. More minor features associated with the disorder include facial dysmorphia, acquired microcephaly, epilepsy, gastrointestinal disturbances, neuropsychiatric diagnoses, and cortical visual impairment. Although rare, premature death due to early stroke or seizure has been reported (*4, 11*). Currently, there is no FDA-approved treatment addressing the underlying mechanism of *HNRNPH2*-NDD, and management of the disorder remains focused on symptomatic treatment (*11*).

In our previous study, we developed and characterized two knockin mouse models of *HNRNPH2*-NDD, one carrying the most prevalent mutation, R206W, and the other carrying the rare P209L mutation. Both models faithfully recapitulated several features of the human clinical syndrome, including reduced survival in male mice, impaired motor and cognitive functions, and increased susceptibility to audiogenic seizures. These phenotypes showed mutation- and dosage-dependent effects, with P209L mice being more severe than R206W mice, and hemizygous males more severely affected than heterozygous females (*12*). In parallel, we characterized an *Hnrnph2* KO mouse, which were phenotypically normal. Interestingly, we found that expression of *Hnrnph1* was significantly increased in *Hnrnph2* KO mice but not in *Hnrnph2* R206W or P209L knockin mice. HNRNPH1 is a paralog of HNRNPH2, with 96% homology at the amino acid level, and the proteins are believed to play similar, and potentially redundant, roles in RNA processing and splicing (*13*). The expression of the two genes is similar in terms of spatial CNS expression, but different with regard to developmental regulation (*12*). Notably, mutations in *HNRNPH1* have been linked to a neurodevelopmental syndrome very similar to *HNRNPH2*-NDD (*14, 15*). Together, these data suggested that increased *Hnrnph1* expression compensates for the loss of *Hnrnph2* in KO mice, likely accounting for the lack of any observable phenotypes. The fact that this increased expression is not observed in *Hnrnph2* knockin mutant mice suggested two possible mechanisms as drivers of disease. The first possibility is a toxic gain of function. Indeed, we observed a decrease in nuclear levels of mutant HNRNPH2 protein in both mouse brains and human cells, with a concomitant increase in cytoplasmic expression. However, the level of mislocalization was modest, with most of the mutant protein remaining in the nucleus. Instead, our data favored an alternative pathological mechanism of complex loss of function of HNRNPH2 driven by failure of HNRNPH1 compensation. Importantly, both of these possible gain-of-function and loss-of-function mechanisms are predicted to respond positively to therapies designed to deplete expression of HNRNPH2 while simultaneously upregulating HNRNPH1. In the current study, we tested this hypothesis by treating mice with a gapmer antisense oligonucleotide (ASO) designed to target murine *Hnrnph2* mRNA in a non-allele-specific manner for RNase H degradation. We found that intracerebroventricular (ICV) injection of the *Hnrnph2* ASO was well tolerated and resulted in dose-dependent decreases in *Hnrnph2* expression and increases in *Hnrnph1* expression in the mouse brain. We further found that both neonatal and juvenile treatment with the ASO rescued or improved multiple phenotypes of *Hnrnph2* mutant P209L and R206W knockin mice. These results provide proof of principle that knockdown of *Hnrnph2* to sufficient levels leads to a compensatory increase in *Hnrnph1* expression, is well tolerated, and is a potential therapeutic strategy for *HNRNPH2*-NDD. Importantly, as the ASO targets both WT and mutant *Hnrnph2*, this strategy may prove effective for *HNRNPH2*-NDD patients regardless of their specific mutation.

## Results

### ASO targeting mouse *Hnrnph2* in a non-allele-specific manner knocks down *Hnrnph2* and upregulates *Hnrnph1*

To develop ASOs that effectively knock down *Hnrnph2* in our mouse model of *HNRNPH2*-NDD, we designed and tested more than 450 gapmer ASOs targeting murine *Hnrnph2* in a non-allele-specific manner for degradation by RNase H. ASOs were initially screened in primary mouse cortical neurons at 20 µM, and the most active ASOs were subsequently evaluated in a 4-point dose-response curve ranging in concentration from

1.2 to 32 µM. The top 19 most efficacious ASO candidates in vitro were subsequently selected for an in vivo 8-week activity and tolerability assessment in WT mice. Briefly, adult C57BL/6J female mice received a single bolus ICV injection of 700 µg *Hnrnph2* ASO or vehicle (PBS) control and were assessed for neurological function and behavior, weight gain, and markers of neuroinflammation, including *Aif1*, IBA1, and CD68 (**table S1**). The ASOs demonstrated variable efficacy, with greater cortical *Hnrnph2* mRNA reduction associated with greater *Hnrnph1* mRNA upregulation (**fig. S1A**). The most potent *Hnrnph2* ASO, ASO1, knocked down *Hnrnph2* mRNA levels by 96% (**fig. S1B**) while increasing *Hnrnph1* mRNA by 63% (**fig. S1C**). ASO1 (hereafter referred to as *Hnrnph2* ASO) also showed a favorable tolerability profile, including no significant effect on markers of neuroinflammation or body weight (**fig. S1, D and E**), and was selected for further studies.

### Potency and duration of action of *Hnrnph2* ASO in mouse brain

To test the potency of the *Hnrnph2* ASO in vivo, we performed ICV injections of WT C57BL/6J pups with a single bolus of 1.875-30 µg of non-targeting control ASO or *Hnrnph2* ASO at postnatal day (PND) 2-4, followed by whole brain harvest at age 3 weeks (**Fig. 1A**). The *Hnrnph2* ASO significantly reduced *Hnrnph2* mRNA levels in a dose-dependent manner as measured by ddRT-PCR, starting with the lowest dose tested (**Fig. 1B**). As previously demonstrated (*12*), depletion of *Hnrnph2* increased *Hnrnph1* mRNA levels in a dose-dependent manner, starting from the second lowest dose tested (**Fig. 1C**). As all doses of control ASO and *Hnrnph2* ASO were well tolerated in this cohort, the highest dose (30 µg) was selected for further neonatal treatment studies.

**Fig. 1.**
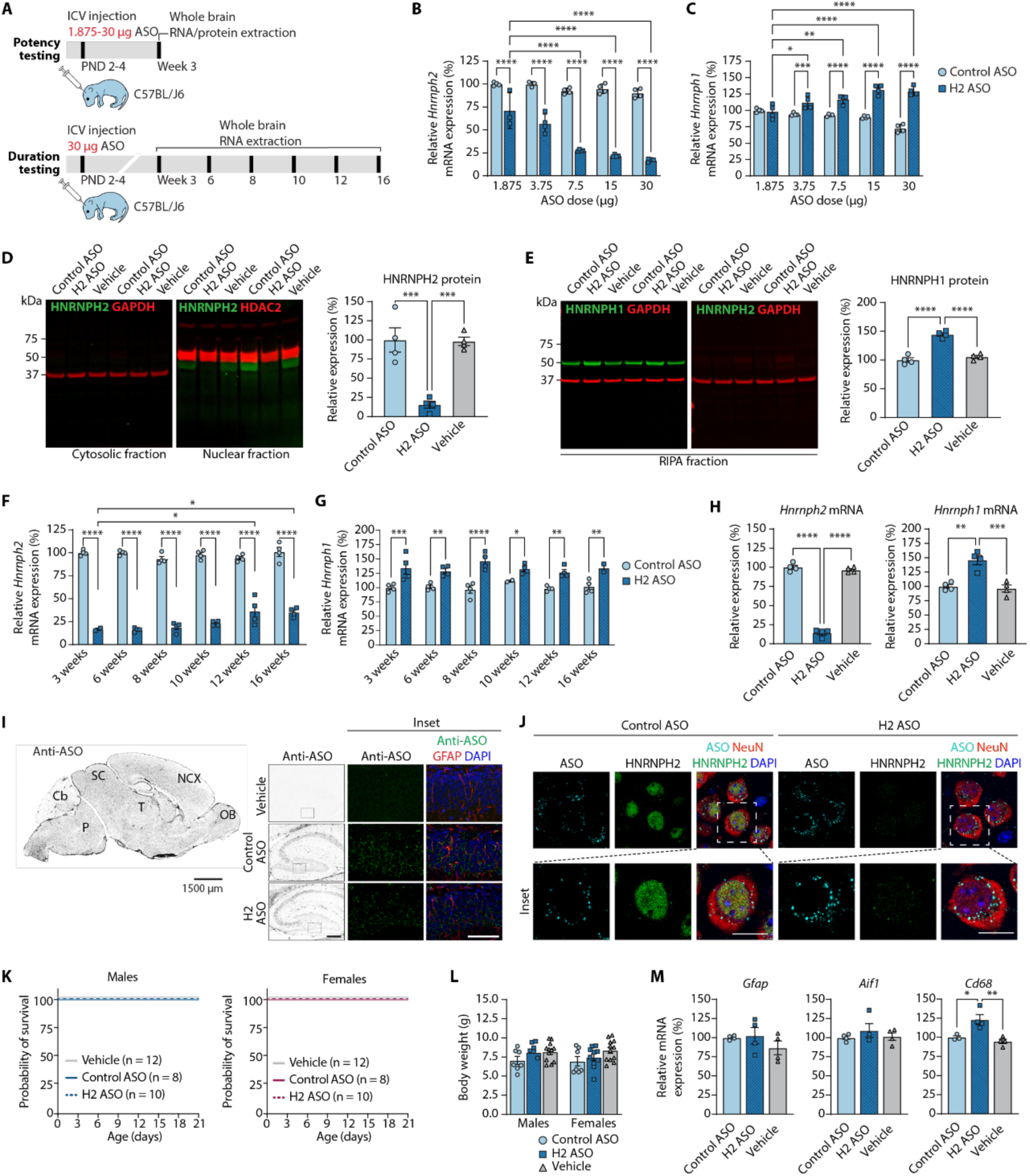
Potency, duration, distribution, and tolerability of neonatal *Hnrnph2* ASO treatment in C57BL/6J mice. (**A**) Design of studies used to generate data in (B-M). All panels show data collected from C57BL/J6 mice at 3 weeks of age (except as indicated in F, G, and K) and with 30 μg dose of ASO (except as indicated in B). (**B-C**) ddRT-PCR of whole brain tissues showing *Hnrnph2* and *Hnrnph1* mRNA levels with increasing doses of ASO. *n* = 3-4 mice per dose. (**D-E**) Immunoblot of whole brain tissues showing HNRNPH2 and HNRNPH1 protein levels in indicated lysate fractions. *n* = 4 mice per treatment. (**F-G**) ddRT-PCR of whole brain tissues showing *Hnrnph2* and *Hnrnph1* mRNA levels at indicated time points. *n* = 2-4 mice per age. (**H**) ddRT-PCR of whole brain tissues showing *Hnrnph2* and *Hnrnph1* mRNA levels in mice treated with ASO or vehicle. *n* = 4 mice per treatment. (**I**) Left, low-magnification image of mouse brain showing ASO distribution (scale bar, 300 μm). OB: Olfactory bulb. NCX: Neocortex. T: Thalamus. SC: superior colliculus. Cb: Cerebellum. P: Pons. Right, higher-magnification images (inset scale bar, 50 μm) of hippocampus. (**J**) Staining of neocortex with antibodies against ASO, HNRNPH2, and the neuronal marker NeuN. Scale bar, 7 μm. (**K**) Kaplan-Meier curves for indicated genotypes and treatments. (**L**) Body weight for indicated genotypes and treatments. *n* = 8-12 mice per treatment. (**M**) ddRT-PCR of whole brain tissues showing *Gfap, Aif1,* and *Cd68* expression. *n* = 3-4 mice per treatment. For all statistical analyses, \*\*\*\**P* < 0.0001, \*\*\**P* < 0.001, \*\**P* < 0.01, \**P* < 0.05. For panels (D), (E), (H), (M): 1-way ANOVA with Tukey’s multiple comparison. For panels (B), (C), (F), (G), (L): 2-way ANOVA with Tukey’s multiple comparison. All bar graphs show data as mean ± SEM.

The changes in *Hnrnph2* and *Hnrnph1* mRNA levels with 30 μg *Hnrnph2* ASO were mirrored by changes in protein expression as measured by Western blot of nuclear and cytoplasmic fractions of whole brain tissue at 3 weeks of age. Due to its low expression levels, HNRNPH2 protein was detectable by an HNRNPH2-specific antibody (*12*) (**fig. S2**) only in nuclear fractions, where it was significantly reduced in *Hnrnph2* ASO-treated samples compared to control ASO- and vehicle-treated samples (**Fig. 1D and fig. S2**). In contrast, HNRNPH1 was readily detected in whole-cell RIPA fractions using an HNRNPH1-specific antibody (**fig. S2**) and showed a significant increase following *Hnrnph2* ASO treatment (**Fig. 1E and fig. S2**), consistent with their respective changes in mRNA levels.

To determine the duration of action of the *Hnrnph2* ASO, we performed a single-bolus ICV injection of 30 µg control ASO or *Hnrnph2* ASO in C57BL/6J pups at PND 2-4, followed by whole brain harvest at 3-16 weeks of age (**Fig. 1A**). We observed significant knockdown of *Hnrnph2* mRNA until the maximum time tested (16 weeks of age), although the level of knockdown began to decline at 12 weeks as compared to 3 weeks (**Fig. 1F**). Similarly, *Hnrnph1* mRNA was significantly increased compared to control ASO treatment up to 16 weeks of age, with no apparent reduction in upregulation at later time points as compared to 3 weeks (**Fig. 1G**). As expected, no significant differences were observed between the 30 µg control ASO and vehicle treatments (**Fig. 1H**). These data demonstrate that the effect of the *Hnrnph2* ASO is stable for at least 12 weeks in mouse brain after a single early postnatal ICV injection.

### Distribution of *Hnrnph2* ASO in mouse brain

To determine the distribution of ASO in the mouse brain, we used immunofluorescent staining with a polyclonal antibody that selectively recognizes the phosphorthioate backbone of ASOs (*16, 17*). This antibody showed broad distribution of the *Hnrnph2* ASO and control ASO across the brain (**Fig. 1I**). Higher magnification revealed a punctate staining pattern in *Hnrnph2* ASO- and control ASO-treated samples, with no staining evident in vehicle-treated brains (**Fig. 1I**). Costaining with specific markers for neurons, oligodendrocytes, microglia, and astrocytes confirmed the presence of ASO in all four major cell types of the CNS (**fig. S3A**). Lastly, triple staining with the anti-ASO antibody, an HNRNPH2-specific antibody, and neuronal markers revealed downregulation of HNRNPH2 signal in neurons of the neocortex (**Fig. 1J**), hippocampus, and cerebellum (**fig. S3, B and C**) that were positive for the *Hnrnph2* ASO. Together, these data demonstrate that a single ICV bolus injection at PND 2-4 results in broad distribution of the *Hnrnph2* ASO in the mouse brain and efficient reduction of HNRNPH2 protein in all major CNS cell types.

### *Hnrnph2* ASO is well tolerated

To assess the safety of neonatal treatment with the *Hnrnph2* ASO, we performed ICV injection of C57BL/6J pups with a single bolus of 30 µg control ASO, *Hnrnph2* ASO, or vehicle at PND 2-4. Pups were followed for survival until 3 weeks of age, at which point they were weighed and whole brains harvested. We found no significant difference in survival or body weight between any of the treatment groups for males and females (**Fig. 1, K and L**). As concerns have been raised regarding neuroinflammation in response to CNS administration of ASOs (*18*), we next examined the expression of reactive gliosis markers by ddRT-PCR using whole brain tissues collected at 3 weeks of age. *Hnrnph2* ASO-treated brains showed no significant difference in mRNA levels of the astrocytic marker *Gfap* or the microglial marker *Aif1* compared to control ASO- or vehicle-treated samples (**Fig. 1M**). Although *Cd68* transcript levels were significantly increased in *Hnrnph2* ASO-treated brains, *Cd68* is more closely related to lysosomal activity than to overall microglial activation, whereas *Aif1* is a more direct marker of microglial activation (*19*). Importantly, histological analyses revealed no overt signs of neuroinflammation or systemic toxicity in brain (**table S2**) or any organs (**table S3**) following administration of either control or *Hnrnph2* ASO. Together, these data indicate that treatment with 30 µg control ASO and *Hnrnph2* ASO in neonatal WT C57BL/6J mice is well tolerated. As there were no significant differences between the control ASO-treated and vehicle-treated groups, all further neonatal treatment studies were conducted with the non-targeting control ASO and *Hnrnph2* ASO only (no vehicle groups) to reduce the number of animals needed to complete the study.

### Neonatal treatment with *Hnrnph2* ASO does not influence the survival and body weight phenotypes of mutant *Hnrnph2* mice

We next evaluated the effect of the *Hnrnph2* ASO in mutant *Hnrnph2* mice. For these and all subsequent experiments, we evaluated phenotypes in at least two genotype/sex groups. Notably, these mouse models have a consistent pattern of phenotype severity across all phenotypes tested: P209L hemizygous male mice are the most severely affected, followed in decreasing severity by R209W hemizygous mice and P209L heterozygous female mice; R206W heterozygous females show no significant phenotype in most tests (*12*). Thus, we designed our experiments to assess the effects of *Hnrnph2* ASO in the most severely affected group that was testable (i.e., P209L hemizygous males in nearly all assays), along with at least one more mildly affected group.

Following the same protocol as was used for WT C57BL/6J mice, we treated *Hnrnph2* P209L hemizygous males and WT littermates with a single bolus ICV injection of 30 µg control ASO or *Hnrnph2* ASO at PND 2-4 and collected whole brains at 3 weeks of age (**Fig. 2A**). *Hnrnph2* mRNA levels were significantly reduced and *Hnrnph1* significantly increased in *Hnrnph2* ASO-treated samples in both WT and P209L mutant mice (**Fig. 2B**), demonstrating that *Hnrnph2* ASO effectively targets both WT and mutant alleles. Next, we treated *Hnrnph2* P209L and R206W mice with *Hnrnph2* ASO using the same regimen and assessed its impact on body weight and survival up to 8 weeks of age. In control ASO-treated male mice, pairwise comparisons indicated that P209L and R206W mice had significant differences in survival compared to WT mice, as previously reported (*12*); however, the statistical significance of these differences was lost after correction for multiple comparisons (**fig. S4, A and B**). In the *Hnrnph2* ASO-treated group, P209L mice showed significantly decreased survival compared to WT mice even after correction for multiple comparisons (**fig. S4A**), whereas in R206W mice there was no significant difference in survival between *Hnrnph2* ASO-treated mutant or WT males, nor was there a significant effect for *Hnrnph2* ASO treatment compared to control ASO treatment in WT or R206W mice (**fig. S4B**). Consistent with our previous report, we found no mutation-dependent differences in survival of heterozygous P209L female mice compared to their WT littermates, regardless of ASO treatment (**fig. S4C**). Together these results demonstrate that neonatal *Hnrnph2* ASO treatment does not rescue reduced survival of *Hnrnph2* mutant mice.

**Fig. 2.**
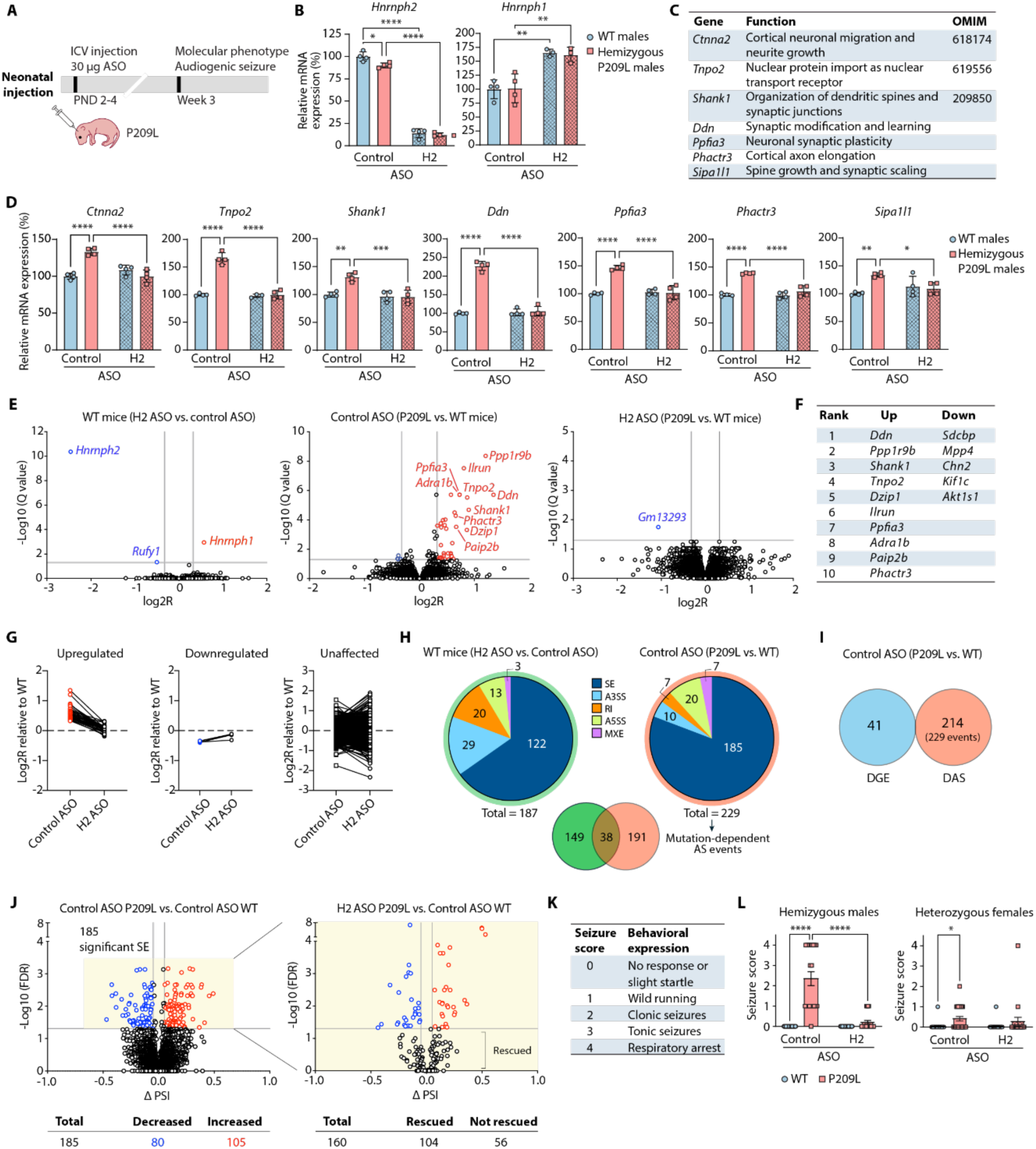
Neonatal treatment with *Hnrnph2* ASO rescues the molecular and audiogenic seizure phenotypes of juvenile *Hnrnph2* P209L mice. (**A**) Design of studies used to generate data in (B-L). All panels show data collected from P209L mutant mice or littermates at 3 weeks of age treated with 30 μg dose of ASO. (**B**) ddRT-PCR of whole brain tissues showing *Hnrnph2* and *Hnrnph1* mRNA levels. (**C**) Seven upregulated genes in both *Hnrnph2* mutant mice and human *HNRNPH2* mutant iPSC-derived neurons. (**D**) ddRT-PCR of whole brain tissues showing expression levels of genes listed in (C). (**E**) Volcano plots showing differentially expressed genes with significantly downregulated (blue) and upregulated (red) genes indicated. (**F**) Top 10 most upregulated and 5 most downregulated genes by log2R from middle plot in (E). (**G**) Effects of *Hnrnph2* ASO on upregulated (left), downregulated (middle), and unchanged (right) genes at the individual gene level. (**H**) RNA-seq-derived differential alternative splicing events grouped by *Hnrnph2* ASO treatment (left) and P209L mutation (right). SE: skipped exon. A3SS: alternative 3′ splice site. RI: retained intron. A5SS: alternative 5′ splice site. MXE: mutually exclusive exon. Overlap between *Hnrnph2* ASO-dependent and P209L mutation-dependent events is shown. (**I**) No overlap between P209L mutation-dependent events identified by differential gene expression (DGE) and differential alternative splicing (DAS) analyses. (**J**) Left, volcano plot showing the 185 significant skipped exon (SE) events in control ASO-treated P209L mutant mice with significantly increased (red) and decreased (blue) events. Right, volcano plot showing the effect of *Hnrnph2* ASO treatment on P209L mutation-dependent skipped exon events. (**K-L**) Audiogenic seizure susceptibility scoring (K) and assessment (L) of indicated mice. For panel B and D, 2-way ANOVA with Sidak’s multiple comparison, *n* = 3-4 mice per treatment. For panel L, non-parametric Scheirer–Ray–Hare test with Mood’s median test, control ASO WT males *n* = 20, control ASO P209L males *n* = 30, *Hnrnph2* ASO WT males *n* = 20, *Hnrnph2* ASO P209L males *n* = 29, control ASO WT females *n* = 26, control ASO P209L females *n* = 27, *Hnrnph2* ASO WT females *n* = 30, *Hnrnph2* ASO P209L females *n* = 19. For all statistical analyses, \*\*\*\**P* < 0.0001, \*\*\**P* < 0.001, \*\**P* < 0.01, \**P* < 0.05. Data shown as mean ± SEM.

Similarly, although control ASO-treated *Hnrnph2* P209L and R206W males, but not P209L females, showed reduced body weight compared to WT littermates at 8 weeks, this phenotype was not rescued by *Hnrnph2* ASO treatment (**fig. S4, D to F**), suggesting that this aspect of the phenotype may be driven by peripheral factors that are not rescued by central administration of ASO. Importantly, there was no significant difference in body weight between control ASO and *Hnrnph2* ASO-treated WT or *Hnrnph2* mutant mice, indicating that neonatal treatment with the *Hnrnph2* ASO is well tolerated in R206W males and P209L mice.

### Neonatal treatment with *Hnrnph2* ASO rescues the molecular phenotype of juvenile mutant *Hnrnph2* mice

Next, we investigated whether the *Hnrnph2* ASO was able to rescue the molecular phenotype of *Hnrnph2* P209L and R206W male mice previously identified by RNA-seq (*12*). Using whole brain samples from *Hnrnph2* P209L and R206W hemizygous males and WT littermates treated with a single bolus ICV injection of 30 µg control ASO or *Hnrnph2* ASO at PND 2-4, we assessed the mRNA levels of seven differentially expressed genes identified in our original study (**Fig. 2C**). These genes (*Ctnna2*, *Tnpo2*, *Shank1*, *Ddn*, *Ppfia3*, *Phactr3*, and *Sipa1l1*) were chosen based on their differential expression in 8-week-old *Hnrnph2* R206W and 3-week-old P209L hemizygous male mice, and in human HNRNPH2 R206W, P209L, and R206Q induced pluripotent stem cell (iPSC)-derived neurons, while not being significantly changed in *Hnrnph2* KO hemizygous male mice or human *HNRNPH2* KO iPSC-derived neurons (*12*). Using ddRT-PCR, we found that neonatal *Hnrnph2* ASO treatment restored the expression of all 7 genes in *Hnrnph2* P209L mutant males to WT levels at 3 weeks of age (**Fig. 2D**). A similar rescue of gene expression was observed in *Hnrnph2* R206W mutant males following neonatal *Hnrnph2* ASO treatment (**fig. S5, A to D**). Notably, in WT mice, *Hnrnph2* ASO treatment did not change the mRNA levels of any of the seven genes tested, likely due to functional compensation by upregulation of *Hnrnph1* in response to reduced *Hnrnph2* (**Fig. 2D and fig. S5D**).

To examine the broader impact of neonatal ASO treatment, we performed RNA-seq analyses on whole brains from 3-week-old *Hnrnph2* P209L and R206W male mice treated with a single bolus ICV injection of 30 µg control ASO or *Hnrnph2* ASO at PND 2-4. In WT mice, only three genes, including *Hnrnph2* and *Hnrnph1,* were differentially expressed following *Hnrnph2* ASO treatment compared to control ASO (**Fig. 2E**, left), supporting our hypothesis of functional compensation by *Hnrnph1* upregulation in response to reduced *Hnrnph2* and consistent with the previously reported absence of phenotypes in *Hnrnph2* KO mice (*12*). Moreover, the lack of significant changes in the expression of other genes following *Hnrnph2* ASO treatment supports the specificity and safety of the ASO. Within the control ASO-treated groups, P209L male mice had 41 differentially expressed genes compared with WT mice, with 36 genes upregulated and only 5 genes downregulated (**Fig. 2E**, middle, and **Fig. 2F**), consistent with a prior report that HNRNPH1/2 primarily act as negative regulators of target gene expression (*20*). Importantly, there was only 1 significantly differentially expressed gene in P209L samples compared to WT samples after *Hnrnph2* ASO treatment (**Fig. 2E**, right). At the individual gene level, the expression of both upregulated and downregulated genes in P209L mice was restored to WT levels with *Hnrnph2* ASO (**Fig. 2G**), indicating that neonatal *Hnrnph2* ASO treatment broadly rescues the molecular phenotypes observed in juvenile *Hnrnph2* P209L mice.

We obtained similar results from juvenile *Hnrnph2* R206W male mice after neonatal *Hnrnph2* ASO treatment. Consistent with their milder behavioral phenotype compared to P209L mutant mice (*12*), R206W mutant mice had only 14 differentially expressed genes compared to WT mice in the control ASO-treated group, with 13 genes significantly upregulated, and only 1 gene downregulated (**fig. S5, E** middle, **and F**). All of these expression changes were restored to WT levels following *Hnrnph2* ASO treatment (**fig. S5G**). We note that *Hnrnph1* transcript levels were not significantly increased in this RNA-seq analysis of samples treated with *Hnrnph2* ASO, although a significant increase was detected by ddRT-PCR assay from the same samples (**fig. S5B**). This discrepancy could be due to the targeted nature and higher sensitivity of ddRT-PCR, which is better able to detect subtle changes in transcript abundance compared to the more global approach of RNA-seq. Taken together, these results demonstrate that neonatal *Hnrnph2* ASO treatment is effective in correcting the gene expression profile of both *Hnrnph2* P209L and R206W mutant juvenile males, bringing the expression of all significantly differentially expressed genes back to WT levels.

Given the important role of HNRNPH1/2 in regulating alternative splicing (*2*), we also analyzed the RNA-seq data for differential alternative splicing (DAS) events. In contrast to differential gene expression (DGE) (**Fig. 2, E to G** and **fig. S5, E to G**), *Hnrnph2* ASO treatment in WT mice resulted in a substantial number of DAS events. This may reflect incomplete compensation for HNRNPH2 specific splicing events by HNRNPH1 and/or the introduction of novel, HNRNPH1-specific splicing events due to its upregulation. The most common DAS event in *Hnrnph2*-treated WT mice was skipped exons (SE), accounting for 65% of all events, with 81 SE events upregulated and 41 downregulated (**Fig. 2H**, left). In the control ASO-treated group, P209L mutant mice also exhibited a substantial number of DAS events compared to WT mice, again with SE events (81%) being the most common (**Fig. 2H**, right). Notably, there was no overlap between differentially expressed genes and differentially spliced genes, suggesting that direct alternative splicing of the identified DGE genes is not the primary mechanism underlying changes in gene expression by the P209L mutation (**Fig. 2I**).

We next assessed whether *Hnrnph2* ASO treatment could rescue the DAS events induced by the P209L mutation by examining individual splicing events. To do this, we focused on SE events, as they represent the majority of DAS events. Control ASO-treated P209L mutant mice had 185 significantly altered SE events compared with control ASO-treated WT mice, including 105 upregulated and 80 downregulated events (**Fig. 2J**). *Hnrnph2* ASO treatment restored 104 of these SE events (∼56%) to levels not significantly different from WT, while 56 events (∼30%) remained differentially spliced (**Fig. 2J**). Twenty-five SE events (∼14%) were either not detected by RNA-seq or filtered out from the final dataset during data analysis due to low read counts or extreme percent spliced in (PSI) values (*21*).

Control ASO-treated R206W mutant male mice exhibited 549 DAS events compared to control ASO-treated WT mice, with SE events comprising the majority (459, ∼84%) (**fig. S5H**), a substantial increase in total DAS events that is likely attributable to batch effects, given a similar increase in control ASO-treated WT mice in this experiment. Only 1 gene, *Stmn4*, was both differentially expressed and differentially spliced (**fig. S5I**). This minimal overlap between DEG genes and DAS genes suggests that direct alternative splicing of the DEG genes is not the primary mechanism underlying changes in gene expression by the R206W mutation. Following *Hnrnph2* ASO treatment, approximately 52% of significant SE events were restored to levels not significantly different from WT, 35% remained differentially spliced, and 14% were not detected or filtered out from the final dataset during data analysis (**fig. S5J**). Together, these data suggest that *Hnrnph2* ASO treatment is effective in rescuing a substantial proportion of the alternative splicing abnormalities caused by both the P209L and R206W mutation, although some splicing defects persist.

### Neonatal treatment with *Hnrnph2* ASO rescues the audiogenic seizure phenotype of juvenile *Hnrnph2* P209L mice

To further investigate the potential therapeutic effects of knocking down *Hnrnph2*, we tested whether neonatal *Hnrnph2* ASO treatment could rescue behavioral deficits associated with mutant mice. In our initial characterization of *Hnrnph2* mutant mice, we identified increased susceptibility to audiogenic seizures as a robust behavioral phenotype. Audiogenic seizures are generalized seizures triggered by exposure to high-intensity sound and serve as a model to study sensory hypersensitivity and sudden unexpected death in epilepsy (SUDEP; (*22, 23*)). In that study, *Hnrnph2* P209L hemizygous males were the most severely affected, followed by *Hnrnph2* R206W hemizygous males, and *Hnrnph2* P209L heterozygous females with a mild phenotype (*12*). *Hnrnph2* R206W heterozygous females did not have a significant increase in audiogenic seizure susceptibility compared to their WT littermates (*12*). To assess the impact of ASO treatment on susceptibility to audiogenic seizure, we treated P209L hemizygous males, P209L heterozygous females, and R206W hemizygous males, as well as their WT littermates, with a single bolus ICV injection of 30 µg control ASO or *Hnrnph2* ASO at PND 2-4. Mice were maintained until 3 weeks of age, at which point they were tested for audiogenic seizure susceptibility (**Fig. 2A**). We found that neonatal *Hnrnph2* ASO treatment rescued the increased audiogenic seizure susceptibility in juvenile *Hnrnph2* P209L hemizygous males, with no statistically significant difference between WT and mutant mice in the *Hnrnph2* ASO-treated group (**Fig. 2 K and L**). The milder phenotype of *Hnrnph2* P209L heterozygous females was also improved following neonatal *Hnrnph2* ASO treatment (**Fig. 2, K and L**). Although the seizure score of control ASO-treated R206W hemizygous males tended to be increased compared to their WT littermates, it failed to reach statistical significance and neonatal *Hnrnph2* ASO treatment had no significant effect (**fig. S5, K** and **L**).

We previously identified an increased incidence of hydrocephalus as a prominent phenotype in *Hnrnph2* mutant mice (*12*). Although there is no evidence of hydrocephalus in patients with *HNRNPH2* mutations, acquired microcephaly is present in approximately half of cases (*4*). Although the two conditions are typically viewed as mutually exclusive, cases of microcephalic hydrocephalus suggest potential overlap in their pathogenesis, including impaired fetal neural stem cell proliferation (*24*). In our mouse models, hydrocephalus exhibited incomplete penetrance and variable presentation: some mice presented with domed heads, whereas others had normal skulls but ventricular enlargement by MRI scan (*12*). Using the same neonatal treatment regimen as mice tested for audiogenic seizure (**Fig. 2A**), we found that *Hnrnph2* ASO treatment did not rescue the increased incidence of hydrocephalus in *Hnrnph2* P209L hemizygous males (**fig. S6A**). *Hnrnph2* P209L heterozygous females did not show an increased incidence of hydrocephalus compared to their WT littermates, regardless of treatment (**fig. S6A**). Although control ASO- and *Hnrnph2* ASO-treated R206W hemizygous males tended to have a higher incidence of hydrocephalus compared to their WT littermates, the difference was not significantly different after correcting for multiple comparisons (**fig. S6B**). As several studies have implicated defects in early brain development in the pathogenesis of congenital hydrocephalus (*25–27*), it is possible that neonatal ASO treatment may be too late to rescue the hydrocephalus phenotype in *Hnrnph2* mutant mice.

### Neonatal treatment with *Hnrnph2* ASO improves motor function of adult *Hnrnph2* P209L male and female mice

We next assessed the effect of neonatal *Hnrnph2* ASO treatment on motor phenotypes we previously identified in *Hnrnph2* mutant knockin mice. To assess the impact of *Hnrnph2* ASO treatment across a range of phenotype severities, we treated P209L hemizygous males (severe phenotype), P209L heterozygous females (mild phenotype), and their WT littermates with 30 µg control or *Hnrnph2* ASO by single bolus ICV injection at PND 2-4. At 8 weeks old, mice were tested for balance and coordination, muscle strength, and gait changes (**Fig. 3A**). In the rotarod, WT mice performed significantly better than P209L mice when treated with control ASO (**Fig. 3B** and **fig. S7A**). However, after *Hnrnph2* ASO treatment, the difference between WT and P209L mice was no longer significant in both males (**Fig. 3B**) and females (**fig. S7A**), suggesting that *Hnrnph2* ASO treatment improves motor deficits in P209L mice. Balance beam performance was also improved in *Hnrnph2* P209L hemizygous males (**Fig. 3C**) and rescued in *Hnrnph2* P209L heterozygous females (**fig. S7B**): whereas control ASO-treated mutants showed increased latency to cross and more hind paw slips compared to WT littermates, *Hnrnph2* ASO treatment reduced hind paw slips to WT levels, but latency to cross was not rescued in *Hnrnph2* P209L hemizygous males (**Fig. 3C**). In females, *Hnrnph2* ASO treatment restored these measures to WT levels (**fig. S7B**). For grip strength, *Hnrnph2* P209L males exhibited slightly weaker muscle strength compared to WT in the control ASO-treated group, though this difference did not reach statistical significance. In *Hnrnph2* ASO-treated males, muscle strength remained impaired (**Fig. 3D**). Of note, we observed that *Hnrnph2* ASO-treated WT males showed significantly higher grip strength compared to control ASO-treated WT males, suggesting a possible treatment effect. Grip strength was improved in *Hnrnph2* ASO-treated *Hnrnph2* P209L females (**fig. S7C**). Finally, reduced stride length was restored in *Hnrnph2* P209L males, with no significant difference between *Hnrnph2* ASO-treated mutants and WT littermates, and a significant increase compared to control ASO-treated mutants (**Fig. 3E**). No significant stride length reduction was observed in *Hnrnph2* P209L females compared to their WT littermates and Hnrnph2 ASO treatment had no effect (**fig. S7D**).

**Fig. 3.**
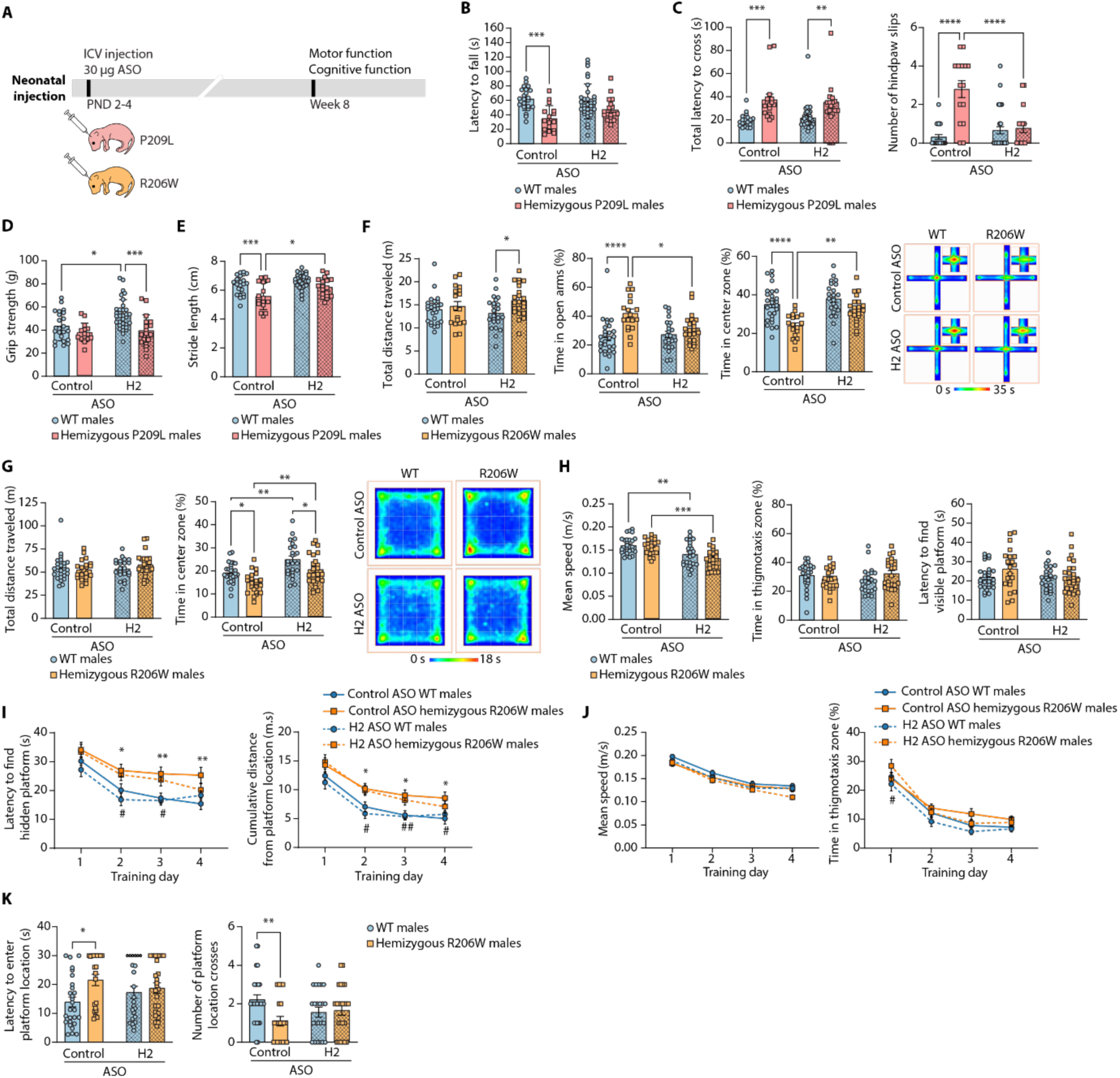
Neonatal treatment with *Hnrnph2* ASO improves motor and cognitive function in adult *Hnrnph2* P209L and R206W male mice. (**A**) Design of studies used to generate data in (B-K). (**B-E**) Effect of neonatal *Hnrnph2* ASO treatment on rotarod performance (B), balance beam performance (C), grip strength (D), and stride length (E) of P209L male mice. (**F-K**) Effect of neonatal *Hnrnph2* ASO treatment on elevated plus maze behavior (F), open field behavior (G), and Morris water maze performance (H-K) in female P209L mice. In (F), heatmaps show the average animals’ center point for the groups for the 5-minute test. In (G), heatmaps show the average animals’ center point for the groups for the 20-minute test. For panels B-H and K, \*\*\*\**P* < 0.0001, \*\*\**P* < 0.001, \*\**P* < 0.01, \**P* < 0.05 by 2-way ANOVA with Sidak’s multiple comparison. For panel I, \*\**P* < 0.01, *\*P* < 0.05 control ASO WT males vs. control ASO R206W males, ##*P* < 0.01, #*P* < 0.05 *Hnrnph2* ASO WT males vs. *Hnrnph2* ASO R206W males, linear mixed effects model with random intercept. Post-hoc comparisons adjusted by FDR within each time point. For (B-E), control ASO WT males *n* = 22, control ASO P209L males *n* = 16, *Hnrnph2* ASO WT males *n* = 33, *Hnrnph2* ASO P209L males *n* = 18. For (G-K), control ASO WT males *n* = 27, control ASO R206W males *n* = 21, *Hnrnph2* ASO WT males *n* = 23, *Hnrnph2* ASO R206W males *n* = 24. All graphs show data as mean ± SEM.

### Neonatal treatment with *Hnrnph2* ASO improves cognitive function in adult *Hnrnph2* R206W male mice

The major and most common feature of *HNRNPH2*-NDD is intellectual disability, while anxiety and other neuropsychiatric symptoms are also often reported and of considerable concern to patients’ parents (*4*). In our original phenotypic characterization of the *Hnrnph2* mouse lines, we found that R206W males have mildly impaired decision making related to approach/avoid conflict and impaired spatial learning and memory (*12*). In that study, we elected not to perform these assays in the *Hnrnph2* P209L males due to their high mortality rate at age 8 weeks as well as concerns regarding their physical ability to participate in a water maze test. Although we did not assess the cognitive phenotype of *Hnrnph2* P209L heterozygous female mice in the original phenotyping study, based on the pattern seen for all other tests, it was predicted to be milder than that of the *Hnrnph2* R206W hemizygous males. To assess the effect of *Hnrnph2* ASO treatment on the impaired decision making and spatial learning and memory previously identified in *Hnrnph2* R206W males and predicted in *Hnrnph2* P209L females, we injected hemizygous R206W males, heterozygous P209L females, and their WT littermates with a single ICV bolus of 30 µg control or *Hnrnph2* ASO at PND 2-4. Mice were maintained until 8 weeks of age, when they were tested first in the elevated plus maze, followed by the open field test, and lastly the Morris water maze (MWM) (**Fig. 3A**).

In the elevated plus maze, total distance traveled, a measure associated with locomotor activity, was not significantly different between WT and R206W male mice when treated with control ASO; however, with *Hnrnph2* ASO treatment, R206W males showed a significant increase compared to WT (**Fig. 3F**). R206W males spent significantly more time in the open arms and less time in the maze center compared to WT mice when treated with control ASO, indicative of impaired decision making related to approach/avoid conflict. In *Hnrnph2* ASO-treated mutants, these parameters were restored to WT levels (**Fig. 3F**). However, interpretation of these results is confounded given the increased general activity in *Hnrnph2* ASO-treated mutants as indicated by increased total distance traveled.

In the open field test, there were no significant differences in total distance traveled, a measure of locomotor activity, between mice regardless of genotype and treatment (**Fig. 3G**). With control ASO treatment, R206W mutant males spent significantly less time in the open field center compared to WT males, suggestive of increased anxiety in the mutants. While *Hnrnph2* ASO-treated mutant mice spent significantly more time in the center zone compared to control ASO-treated mutant mice, the interpretation of this result is confounded by the finding that treatment with the *Hnrnph2* ASO resulted in increased time in the center zone in both WT and mutant males compared to their control ASO counterparts (**Fig. 3G**).

We next assessed spatial learning and memory in the MWM. To ensure that the mice possessed the basic sensory-motor functions and visual ability required to complete the task, we first performed a cued version of the MWM. Swim speed, a parameter associated with locomotion, was significantly reduced in *Hnrnph2* ASO-treated male mice, regardless of genotype (**Fig. 3H**). However, percent time spent in the thigmotaxis zone of the pool, which is associated with anxiety, and latency to find the visible platform were similar in all groups (**Fig. 3H**), demonstrating that ASO treatment did not alter the basic abilities, strategies, and motivation that is required for completing the task. In the training phase of the MWM, which assesses spatial learning, control ASO-treated R206W males showed a significant increase in latency to find the hidden platform and cumulative distance from the hidden platform compared with their WT littermates on days 2, 3, and 4 (**Fig. 3I**), despite similar swim speed and percent thigmotaxis time (**Fig. 3J**), suggesting a mild deficit in spatial learning. *Hnrnph2* ASO treatment did not significantly improve this deficit (**Fig. 3I**). In the probe trial of the MWM to assess memory retention, control ASO-treated R206W males showed a significant increase in latency to first enter the platform location and a significant reduction in the number of platform location crossings compared to their WT littermates, suggesting a deficit in spatial memory (**Fig. 3K**). This deficit was improved by *Hnrnph2* ASO treatment, with no significant difference in either parameter between *Hnrnph2* ASO-treated R206W males and WT littermates (**Fig. 3K**), suggesting that although *Hnrnph2* ASO treatment did not rescue all parameters in the MWM test, it was effective in restoring spatial memory in R206W mutant male mice.

P209L heterozygous females exhibited minimal cognitive or behavioral deficits (**figs. S7, E to I**). The only significant difference between WT and P209L heterozygous females was increased time spent in the open arms of the elevated plus maze (**fig. S7E**), indicative of increased anxiety-like behavior. This phenotype was improved by *Hnrnph2* ASO treatment, with treated mice showing open arm times comparable to WT controls (**fig. S7E**). *Hnrnph2* ASO treatment had effects in other tests that did not show any mutation-dependent phenotypes, including increases in the total distance traveled and time in center zone in the open field test (**fig. S7F**), decreased swim speed and increased latency to find the platform in the MWM cued phase (**fig. S7G**), and isolated effects in day 4 of the MWM training phase (**fig. S7H**).

### Juvenile treatment with *Hnrnph2* ASO rescues molecular and audiogenic seizure phenotypes and improves the motor phenotype of adult *Hnrnph2* P209L mice

We next tested an extended therapeutic window of *Hnrnph2* ASO treatment. To do so, we first tested the potency, duration of action, and tolerability of a juvenile ASO treatment protocol in WT C57BL/6J mice. Mice received a single bolus ICV injection of 30-200 µg of non-targeting control ASO or *Hnrnph2* ASO at PND 27-29, followed by whole brain harvest at age 8 weeks (**Fig. 4A**). Similar to results obtained from our neonatal (PND 2-4) *Hnrnph2* ASO treatment protocol, mice treated with *Hnrnph2* ASO at a juvenile stage showed significantly reduced *Hnrnph2* mRNA levels and increased *Hnrnph1* mRNA levels in a dose-dependent manner (**Fig. 4B and 4C**). This knockdown of *Hnrnph2* mRNA and upregulation of *Hnrnph1* mRNA persisted until the maximum time tested (16 weeks of age) (**Fig. 4D and 4E**). No significant differences were observed between 200 µg control ASO and vehicle treatments (**Fig. 4F**). Similar to the findings with neonatal treatment, juvenile ASO-treated mice showed upregulated *Cd68* transcript levels; however, assessment of other reactive gliosis markers revealed no apparent neuroinflammation following CNS administration of ASOs at this stage (**Fig. 4G**), consistent with results observed in neonatally treated mice. As there were no significant differences between the control ASO-treated and vehicle-treated groups, all further juvenile treatment studies were conducted with the non-targeting control ASO only (no vehicle groups) to reduce the number of animals needed for the study.

**Fig. 4.**
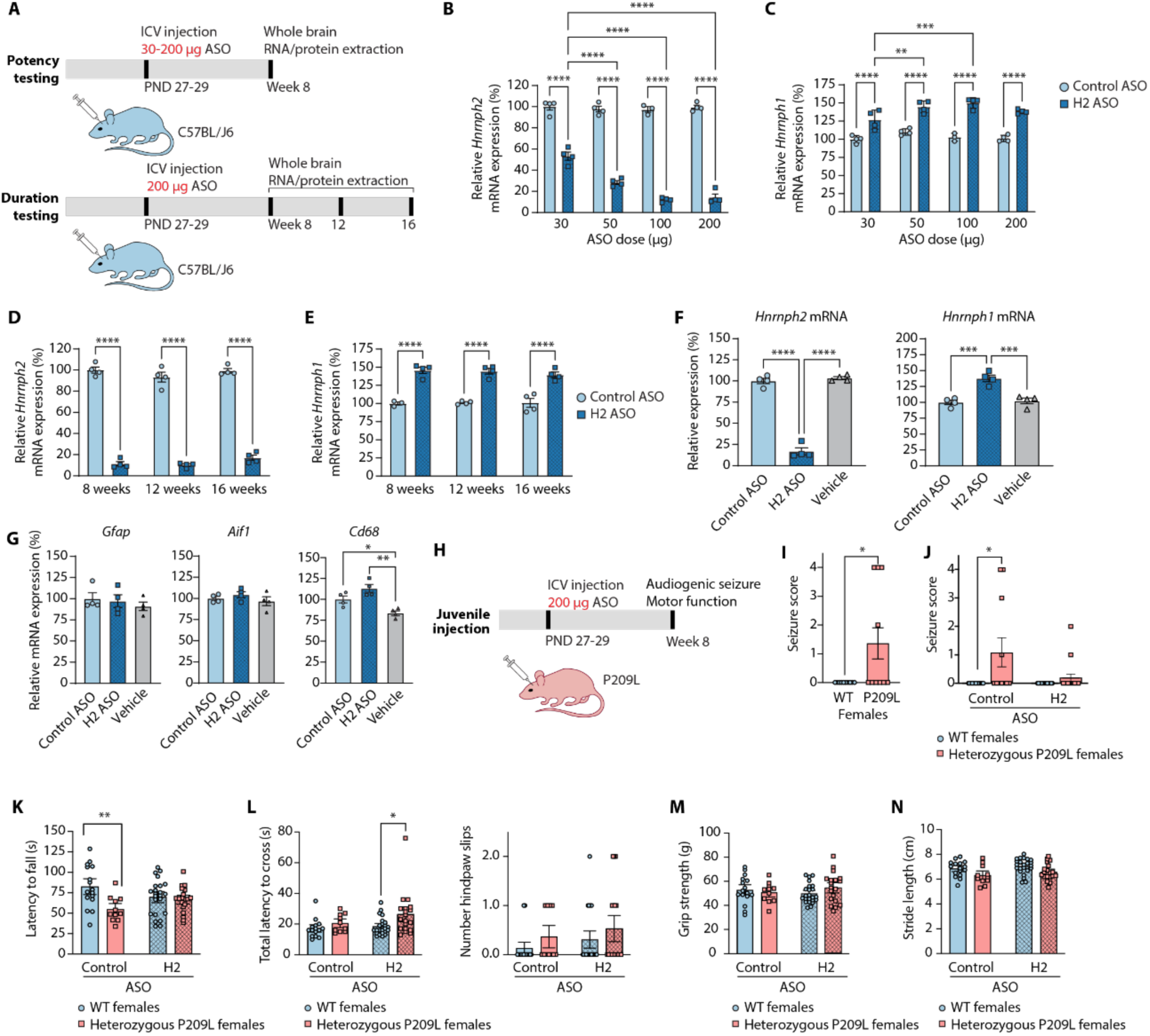
Juvenile treatment with the *Hnrnph2* ASO rescues the audiogenic seizure and motor phenotype of *Hnrnph2* P209L mice. (**A**) Design of studies used to generate data in (B-G). In (B-G), all panels show data collected from C57BL/J6 mice at 8 weeks of age (except as indicated in D) and with 200 μg dose of ASO (except as indicated in B). (**B-C**) ddRT-PCR of whole brain tissues showing *Hnrnph2* and *Hnrnph1* mRNA levels with increasing doses of ASO. *n* = 3-4 mice per dose. (**D-E**) ddRT-PCR of whole brain tissues showing *Hnrnph2* and *Hnrnph1* mRNA levels at indicated time points. *n* = 2-4 mice per age. (**F**) ddRT-PCR of whole brain tissues showing *Hnrnph2* and *Hnrnph1* mRNA levels in mice treated with ASO or vehicle. *n* = 4 mice per treatment. (**G**) ddRT-PCR of whole brain tissues showing *Gfap, Aif1,* and *Cd68* expression. *n* = 3-4 mice per treatment. (**H**) Design of studies used to generate data in (I-N). (**I-J**) Audiogenic seizure susceptibility in untreated (I) and ASO-treated (J) female mice. For (I): WT females *n* = 10, P209L females *n* = 11. For (J): control ASO WT females *n* = 17, control ASO P209L females *n* = 11, *Hnrnph2* ASO WT females *n* = 16, *Hnrnph2* ASO P209L females *n* = 16. (**K-L**) Effect of juvenile ASO treatment on rotarod (K) and balance beam (L) performance. For (K): control ASO WT females *n* = 16, control ASO P209L females *n* = 11, *Hnrnph2* ASO WT females *n* = 23, *Hnrnph2* ASO P209L females *n* = 19. For (L): control ASO WT females *n* = 15, control ASO P209L females *n* = 10, *Hnrnph2* ASO WT females *n* = 23, *Hnrnph2* ASO P209L females *n* = 19. (**M-N**) Effect of juvenile ASO treatment on grip strength (M) and stride length (N) of female mice. Control ASO WT females *n* = 16, control ASO P209L females *n* = 11, *Hnrnph2* ASO WT females *n* = 23, *Hnrnph2* ASO P209L females *n* = 19. All graphs show data as mean ± SEM. For all statistical analyses, \*\*\*\**P* < 0.0001, \*\*\**P* < 0.001, \*\**P* < 0.01, \**P* < 0.05 by 2-way ANOVA with Tukey’s multiple comparison (B-E), 1-way ANOVA with Tukey’s multiple comparison (F-G), non-parametric Mann-Whitney U test (I), non-parametric Scheirer–Ray–Hare test with Mood’s median test (J), or 2-way ANOVA with Sidak’s multiple comparison (K-N).

To assess the effect of juvenile administration of *Hnrnph2* ASO on audiogenic seizure susceptibility and motor function, we treated P209L hemizygous males, P209L heterozygous females, and their WT littermates with 200 µg control ASO or *Hnrnph2* ASO by single bolus ICV injection at PND 27-29. Mice were followed until 8 weeks of age, at which point they were weighed and tested for either audiogenic seizure susceptibility or motor function (**Fig. 4H**). In this larger cohort, several mice suffered seizures immediately after ICV injection, some of whom died as a result, while others recovered. For *Hnrnph2* P209L males and WT littermates, there was no significant effect on the incidence of post-ICV injection seizures for genotype or treatment after pairwise comparisons between groups (**fig. S8A**). Similar results were obtained for *Hnrnph2* P209L females and WT littermates; however, after correction for multiple comparisons, control ASO-treated female mice were significantly more likely to experience post-ICV seizures, with P209L female mutants being slightly more susceptible (**fig. S8B**). Thus, juvenile ASO treatment may induce post-ICV injection seizures, and in female mice, the incidence was significantly increased with control ASO treatment as compared to *Hnrnph2* ASO treatment. To minimize potential confounding effects of acute ASO toxicity on the phenotypes being evaluated, we excluded all mice that experienced post-ICV seizures from subsequent behavioral analyses.

Like neonatal treatment, juvenile treatment resulted in significantly reduced whole brain *Hnrnph2* mRNA levels and significantly increased *Hnrnph1* mRNA levels in *Hnrnph2* ASO-treated samples in both WT and P209L mutant male mice (**fig. S8C**), confirming that the *Hnrnph2* ASO effectively targets both WT and mutant alleles. Importantly, all seven genes previously identified as differentially expressed in *Hnrnph2* P209L mutant males were restored to WT levels following juvenile *Hnrnph2* ASO treatment at 8 weeks of age, while no changes were observed in WT mice (**fig. S8D**). RNA-seq analysis of whole brains from 8-week-old *Hnrnph2* P209L male mice treated with either control or *Hnrnph2* ASO further confirmed these findings. In WT mice, *Hnrnph2* was the only differentially expressed gene following *Hnrnph2* ASO treatment, with no significant increase in *Hnrnph1* transcript levels by RNA-seq (**fig. S8E,** left), although ddRT-PCR from the same samples detected its upregulation (**fig. S8C**). These results support the specificity and safety of the *Hnrnph2* ASO in both neonatal and juvenile stages.

In the control ASO-treated groups, adult P209L male mice exhibited 42 differentially expressed genes compared with WT mice, predominantly upregulated (**fig. S8E**, middle, and **fig. S8F**), mirroring the DGE patterns observed in younger mice. Following *Hnrnph2* ASO treatment, no significant differences in gene expression remained between P209L and WT samples (**fig. S8E**, right), with both upregulated and downregulated genes returning to WT levels (**fig. S8G**).

Analysis of DAS events revealed a substantial number of splicing changes in WT mice treated with *Hnrnph2* ASO, consistent with neonatal treatment. The majority were SE events (64%), with 69 upregulated and 54 downregulated (**fig. S8H**). Control ASO-treated P209L mutant mice also exhibited numerous DAS events compared to WT mice, again primarily SE events (64%) (**fig. S8H**). Minimal overlap was observed between differentially expressed genes and differentially spliced genes, suggesting that changes in alternative splicing do not directly underlie the observed gene expression differences (**fig. S8I**).

To evaluate the rescue of splicing abnormalities, we focused on SE events. Control ASO-treated P209L mutant mice had 91 significantly altered SE events compared with WT mice, including 60 upregulated and 31 downregulated events (**fig. S8J**). Juvenile *Hnrnph2* ASO treatment restored 54 SE events (∼59%) to WT levels, while 17 events (∼19%) remained altered (**fig. S8J**); 20 events (∼22%) were either not detected by RNA-seq or filtered out from the final dataset during data analysis due to low read counts or extreme PSI values (*21*). Together, these data demonstrate that juvenile *Hnrnph2* ASO treatment, like neonatal treatment, is effective in correcting both gene expression and alternative splicing abnormalities caused by the P209L mutation, although some splicing defects persist.

Next, we assessed whether 8-week-old mice exhibit audiogenic seizure susceptibility similar to that observed in 3-week-old mice. Both *Hnrnph2* P209L hemizygous males (**fig. S8K**) and heterozygous females (**Fig. 4I**) showed a significant increase in audiogenic seizure susceptibility compared to their WT littermates. Similar to the 3-week-old male mice that received neonatal treatment, juvenile *Hnrnph2* ASO treatment rescued this increased susceptibility to audiogenic seizures in 8-week-old *Hnrnph2* P209L hemizygous males (**fig. S8L**) and improved it in hemizygous females (**Fig. 4J**).

We next assessed whether juvenile *Hnrnph2* ASO treatment can improve motor function. Due to the reduced survival of *Hnrnph2* P209L hemizygous male mice at age 8 weeks and the large number of mice required for behavioral assays, we elected to perform this experiment in *Hnrnph2* P209L heterozygous female mice. We found that the latency to fall from the rotarod in adult P209L females was improved by juvenile *Hnrnph2* ASO treatment (**Fig. 4K**). In contrast to neonatal control ASO-treated P209L females, juvenile control ASO-treated P209L females did not have a significant deficit in balance beam performance compared to their WT littermates. *Hnrnph2* ASO treatment slightly but significantly increased total latency to cross, but not number of hind paw slips (**Fig. 4L**), in P209L females compared to their WT littermates. Finally, in contrast to neonatal control ASO-treated P209L females, juvenile control ASO-treated P209L females did not show a significant reduction in grip strength or stride length, and this was not changed by *Hnrnph2* ASO treatment (**Fig. 4, M and N**).

### HNRNPH2 protein regulates the expression of *HNRNPH1* by modulating its splicing

In our previous study, we showed that *Hnrnph1* total mRNA levels were significantly increased in the cortex of male *Hnrnph2* KO mice, but not in male *Hnrnph2* R206W or R209L mice. We also identified changes in alternative splicing in the brains of these mice (*12*). Upon further investigation of the RNA-seq data from that study, we found that alternative splicing of *Hnrnph1* was significantly different in *Hnrnph2* KO brains, but not in *Hnrnph2* P209L or R206W brains. Specifically, compared to their WT littermates, *Hnrnph2* KO mice, but not *Hnrnph2* R206W or R209L mice, had decreased skipping of an *Hnrnph1* exon (**Fig. 5A and fig. S9A**). Several studies have identified this exon (exon 5 in murine transcript *Hnrnph1-*202 and human transcript *HNRNPH1-*203) and surrounding introns to be involved in regulation of *Hnrnph1* expression. *HNRNPH1* exon 5 is an essential exon, which, if skipped, generates a frameshift and premature termination codon in a downstream exon (**Fig. 5B**), resulting in nonsense-mediated decay (NMD) of the resulting transcript (*28, 29*). Using a primer set that distinguishes between *Hnrnph1* transcripts including and excluding exon 5 (**fig. S9B**), we found that primary cortical neurons from *Hnrnph2* KO mice express higher levels of the exon 5-including transcript (314 bp) together with lower levels of the exon 5-excluding transcript (175 bp) compared to those from WT mice (**fig. S9, C and D**). Additionally, the non-productive, exon 5-skipped transcript (175 bp) increased in both WT and *Hnrnph2* KO cells treated with the NMD inhibitor cycloheximide (CHX), confirming that this transcript is primarily degraded via NMD (**fig. S9, C and D**).

**Fig. 5.**
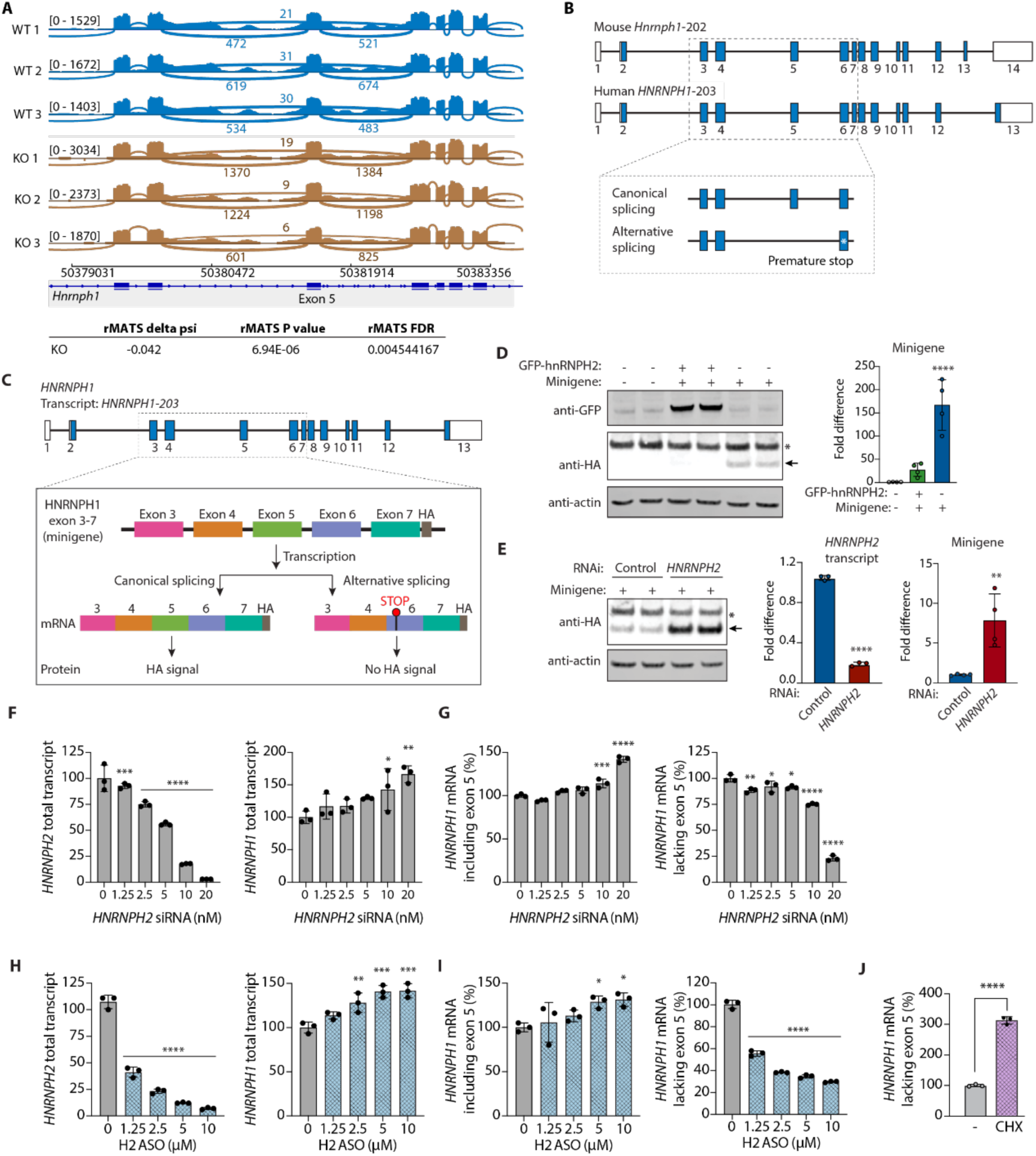
HNRNPH2 protein regulates the expression of *HNRNPH1* by modulating its splicing in human cells and iPSC-derived neurons. (**A**) Differential alternative splicing of *Hnrnph1* in cortex of *Hnrnph2* KO mice. (**B**) Representation of exon/intron organization in *Hnrnph1* and *HNRNPH1*. (**C**) Schematic outputs from the minigene reporter where differential splicing results in inclusion or exclusion of *HNRNPH1* exon 5. The out-of-frame transcript (right) results in the introduction of a premature termination codon in exon 6 and translation of a truncated peptide. (**D**) Left, immunoblot showing HA expression (representing productive splicing) in the presence or absence of GFP-HNRNPH2 overexpression in HEK239T cells. Arrow indicates HA-positive minigene expression; asterisk indicates a nonspecific band. Right, quantification of expression via densitometry. (**E**) Left, immunoblot showing HA expression (representing productive splicing) in presence or absence of RNAi targeting endogenous *HNRNPH2*. Arrow indicates HA-positive minigene expression; asterisk indicates a nonspecific band. (**F**) ddRT-PCR showing *HNRNPH2* and *HNRNPH1* total transcript levels in HEK293T cells after *HNRNPH2* siRNA knockdown. (**G**) ddRT-PCR showing levels of *HNRNPH1* mRNA including or lacking exon 5 in HEK293T cells after *HNRNPH2* siRNA knockdown. (**H**) ddRT-PCR showing *HNRNPH2* and *HNRNPH1* total transcript levels in WT human iPSC-derived neurons after *HNRNPH2* ASO treatment. (**I**) ddRT-PCR showing levels of *HNRNPH1* mRNA including or lacking exon 5 in WT human iPSC-derived neurons after *HNRNPH2* ASO treatment. (**J**) Levels of *HNRNPH1* mRNA lacking exon 5 in WT human iPSC-derived neurons after treatment with cycloheximide. Data are shown as mean ± SD for all panels. For all statistical analyses, \*\*\*\**P* < 0.0001, \*\*\**P* < 0.001, \*\**P* < 0.01, \**P* < 0.05 by student t-test (E and J) or ordinary one-way ANOVA with Dunnett’s multiple comparison (D, F-I).

We next examined the effect of *Hnrnph2* mutations. In control ASO-treated mouse brains, there was no significant difference in the level of *Hnrnph1* non-productive splicing between WT and *Hnrnph2* P209L as measured by ddRT-PCR, mirroring the previously published RNA-seq results (**fig. S9A**). *Hnrnph2* ASO treatment, however, decreased this non-productive splicing in both WT and P209L mice (**fig. S9E**). Together, these results show that *Hnrnph2* KO or significant knockdown by *Hnrnph2* ASO, but not P209L mutation alone, results in a decreased percentage of non-productive splicing of *Hnrnph1*.

The *HNRNPH1* essential exon is flanked by detained introns, a class of retained introns that have been proposed to serve as a pool of nucleus-detained, polyadenylated mRNA for the cell to quickly and efficiently fine-tune gene expression to respond to environmental stimuli and stress (*30, 31*). In Mantle cell lymphoma, mutations in the intron surrounding the essential exon of *HNRNPH1* disrupt binding of HNRNPH1 protein to intronic motifs and increase the inclusion of this exon, leading to increased HNRNPH1 protein expression and inferior clinical outcomes (*32*). Given the high homology between HNRNPH1 and HNRNPH2 proteins and the emerging theme of self- and cross-regulation among RNA-binding protein paralogs through modulation of unproductive splicing (*33–35*), we hypothesized that HNRNPH2 might regulate the expression of *HNRNPH1* by promoting the skipping of its essential exon. To test this, we expressed a minigene containing the genomic sequence for *HNRNPH1* from exons 3 through 7, including all intronic sequences (**Fig. 5C**) (*32*) in HEK293T cells, and examined the effect of overexpressing HNRNPH2 protein on the HA-tagged HNRNPH1 peptide. We found that expression of GFP-HNRNPH2 significantly reduced expression of HA-HNRNPH1, reflecting increased non-productive splicing (i.e., essential exon skipping) (**Fig. 5D**, arrow). Additionally, GFP-HNRNPH2 overexpression was associated with a marked trend toward reduced endogenous HNRNPH1 protein levels (**fig. S9F**). Conversely, inhibiting endogenous *HNRNPH2* expression by RNAi promoted productive splicing (i.e., essential exon inclusion), as reflected by increased levels of the HA-tagged peptide (**Fig. 5E**). This effect was also observed at the endogenous *HNRNPH1* transcript level: RNAi-mediated KD of *HNRNPH2* led to higher total *HNRNPH1* transcript levels (**Fig. 5F**) accompanied by increased inclusion and decreased skipping of exon 5 in a dose-dependent manner (**Fig. 5G**). Together, these data suggest that HNRNPH2 protein, like HNRNPH1 protein, regulates the expression of *HNRNPH1* by modulating its splicing, thus providing molecular insight into how the *Hnrnph2* ASO upregulates *Hnrnph1* expression.

### Research ASO targeting human *HNRNPH2* knocks down *HNRNPH2* and upregulates *HNRNPH1* likely by modulating its alternative splicing

To test whether the compensatory upregulation of *Hnrnph1* in response to *Hnrnph2* ASO treatment observed in the mouse brain is conserved in human neuronal cells, we treated human WT iPSC-derived neurons with a research ASO targeting human *HNRNPH2* for degradation by RNase H (**fig. S9, G and H**). Whereas control ASO had no effect (**fig. S9I**), *HNRNPH2* ASO treatment from day 7 to 21 of neuronal maturation resulted in a significant knockdown of *HNRNPH2* together with increased total *HNRNPH1* mRNA levels in a dose-dependent manner (**Fig. 5H**). This increase in *HNRNPH1* mRNA was also reflected at the protein level, with endogenous HNRNPH1 protein levels increasing following HNRNPH2 ASO treatment (**fig. S9J**). Digital droplet RT-PCR with splicing-specific assays (*32*) showed increased inclusion and decreased skipping of exon 5 in *HNRNPH1* transcripts after *HNRNPH2* ASO treatment (**Fig. 5I**). Since exon 5-skipped transcripts are subject to NMD, as evidenced by a threefold increase after CHX treatment (**Fig. 5J**), the true level of nonproductive transcripts generated is conceivably higher. In summary, our data suggest that the compensatory upregulation of *HNRNPH1* upon *HNRNPH2* ASO treatment is conserved in human cells and that HNRNPH2 modulation of alternative splicing of *HNRNPH1* is likely involved in the mechanism of compensation.

## Discussion

*HNRNPH2*-NDD has a significant impact on the quality of life of patients and their families. Patients typically develop symptoms early in life, including infant feeding difficulties and failure to thrive. Major features of the disorder include developmental delay/intellectual disability, motor and language impairments, and growth and musculoskeletal abnormalities. Facial dysmorphia, acquired microcephaly, behavioral and psychiatric disorders, epilepsy, gastrointestinal disturbances, and cortical visual impairment are also common, though not present in all patients (*36*). Variation in the clinical spectrum is speculated to result from location of the pathogenic mutation in the HNRNPH2 protein (*6*), skewed X chromosome inactivation in the case of females (*5*), and sex (*4*). Finally, although rare, early death due to stroke or seizures has been reported (*4*), highlighting an urgent need for a treatment targeting the root cause of the disease.

Our previous study characterizing *Hnrnph2* KO and mutant knockin mice revealed a mechanism for genetic compensation by *Hnrnph1*. Whereas homologous knockin of human HNRNPH2 R206W or P209L mutations into the mouse *Hnrnph2* gene resulted in a highly penetrant, disease-relevant phenotype, KO of *Hnrnph2* was well tolerated. Loss of HNRNPH2 protein in KO mice triggered upregulation of *Hnrnph1*, the autosomal conserved paralog of *Hnrnph2*. In contrast, modestly reduced levels of mutant HNRNPH2 protein in the nucleus due to impaired nuclear import failed to trigger *Hnrnph1* upregulation. Together, these data suggested either a toxic gain-of-function mechanism for mutant cytoplasmic *Hnrnph2* or a complex loss-of-function mechanism with failed *Hnrnph1* compensation. We hypothesized that both mechanisms would respond positively to a therapeutic strategy designed to deplete HNRNPH2 protein while also triggering compensatory upregulation of *HNRNPH1*.

In the current study, we designed and tested gapmer ASOs targeting *Hnrnph2* in a non-allele-specific manner in a mouse model of *HNRNPH2*-NDD. We found that a single ICV injection of our lead ASO produced long-lasting reductions in *Hnrnph2* expression in whole brain on both the mRNA and protein level accompanied by compensatory increases in the levels of *Hnrnph1* mRNA and total HNRNPH1 protein. The ASO showed widespread distribution across the mouse brain and was well tolerated, with no significant effect on survival, body weight, or signs of neuroinflammation. Both neonatal and juvenile treatments with *Hnrnph2* ASO restored the expression of differentially expressed genes to WT levels and to a lesser extent corrected differentially spliced genes. Importantly, RNA-seq analyses consistently identified seven commonly upregulated genes that we had previously characterized, and these genes were restored to WT levels by *Hnrnph2* ASO treatment in both *Hnrnph2* P209L and R206W mice. These findings suggest that *Hnrnph2* ASO treatment effectively rescues key molecular phenotypes associated with these mutations. Strikingly, both neonatal and juvenile treatment paradigms completely rescued the severe audiogenic seizure susceptibility of *Hnrnph2* P209L male mice and improved the milder seizure phenotype of *Hnrnph2* P209L females. Furthermore, some facets of motor function were also rescued or improved by neonatal or juvenile *Hnrnph2* ASO treatment in adult *Hnrnph2* P209L male and female mice. We also found that neonatal treatment with *Hnrnph2* ASO rescued the anxiety and/or impaired decision-making phenotype of hemizygous R206W males and heterozygous P209L females and significantly improved spatial memory of R206W males. Unfortunately, due to their high mortality rate at age 8 weeks, as well as concerns regarding their physical ability to perform the tasks, we were unable to perform these tests on hemizygous P209L males. Together, these findings suggest that ASO-mediated *HNRNPH2* knockdown could offer a promising therapeutic approach for *HNRNPH2-*NDD patients.

Notably, the body weight, survival, and hydrocephalus phenotypes present in some *Hnrnph2* mutant mice as reported in our previous paper (*12*) were not rescued by *Hnrnph2* ASO treatment. Although reduced life expectancy is not an established feature of the disorder, premature death due to early stroke or seizure has been reported in patients (*4, 11*) and is of major concern to families. In contrast, more than half of patients are reported to have experienced difficulty with feeding and weight gain (*4*). Although we have not identified the specific cause of reduced body weight and survival in *Hnrnph2* mutant mice, it is conceivable that these phenotypes could be driven, at least in part, by peripherally expressed mutant *Hnrnph2* that is not rescued by central administration of the ASO. An ASO that is administered peripherally may be able to treat these and other peripheral symptoms and should be considered as an alternative or adjunctive treatment strategy. Similarly, the exact cause of the increased incidence of hydrocephalus in our *Hnrnph2* mutant mice is unknown, although we did rule out aqueduct blockage in our original phenotyping study. It is possible that the observed hydrocephalus resulted from changes in early brain development (*25, 26*) that occurred prior to ASO administration and were not reversible. Therefore, symptoms that originate prenatally or early postnatally such as hydrocephalus may not be responsive to postnatal treatment and this should be recognized as a limitation of the proposed postnatal ASO-mediated treatment of *HNRNPH2*-NDD.

Although most individuals with *HNRNPH2*-NDD present with symptoms before the age of 12 months (*11*), it typically takes several years to obtain a genetic diagnosis. Therefore, unless widespread newborn screening becomes available, any future disease-altering treatment would likely be administered well after the newborn stage. This is a concern for potential disease-altering treatments of NDDs, as the early childhood period is characterized by the greatest neuroplasticity and is therefore the most beneficial time for intervention (*37*). We are encouraged, however, by the developmental expression profile of *HNRNPH2*: both the mouse and human gene appear to be expressed at relatively low but consistent levels throughout development, from early embryonic stages to adulthood. In contrast, *HNRNPH1* is most highly expressed during early embryonic stages, declining during later fetal and neonatal stages before stabilizing during late infancy into adulthood (*12, 38*). This pattern suggests that *HNRNPH1* may govern early brain developmental processes that are gradually shared with *HNRNPH2* at later and/or post-developmental stages and raises the possibility that treatment targeting *HNRNPH2* mutations may prove to be beneficial even in later developmental stages.

While the time scales characterizing human and altricial rodent brain development differ substantially, the sequence of key events are remarkably conserved. Rodents are born with a relatively immature CNS and experience greater neurogenesis with accelerated brain development and growth postnatally. In contrast, the brains of human neonates are much more developed, although certain processes such as neocortical myelination and synaptic maturation extend well into adulthood (*39*). Thus, although it remains unclear exactly how the mouse neurodevelopmental stages translate to the clinical treatment window for *HNRNPH2*-NDD, our juvenile ASO treatment data suggest that patients may benefit from this therapeutic strategy even after the newborn stage.

In summary, this preclinical study demonstrates the therapeutic potential of non-allele-specific ASO-mediated knockdown of *HNRNPH2* and suggests that this approach may be well tolerated in patients. Advancing beyond these preclinical proof-of-concept studies will require detailed toxicological assessments and further optimization of ASO chemistry and delivery to ensure both efficacy and safety. Reflecting this progress, an ASO-based clinical trial in pediatric *HNRNPH2*-NDD patients was recently launched based on this work. Importantly, this approach may be of benefit to all *HNRNPH2*-NDD patients, regardless of their specific mutation.

## Materials and Methods

### Study design

In this study, we tested whether an ASO that depletes murine *Hnrnph2* while at the same time upregulating *Hnrnph1* can rescue phenotypes of *Hnrnph2* mutant P209L and R206W knockin mice, thereby setting the stage for an ASO-based treatment approach for *HNRNPH2*-NDD. Murine *Hnrnph2* ASOs were first screened in vitro in primary mouse cortical neurons for efficacy in knocking down *Hnrnph2* and upregulating *Hnrnph1*, followed by in vivo safety assessment in adult WT mice. Subsequent in vivo experiments on potency, duration of action, and distribution of the lead ASO were performed in WT mice, followed by testing for effects on molecular and behavioral phenotypes of *Hnrnph2* mutant mice. Both neonatal (PND 2-4) and juvenile (PND 27-29) treatments were tested with the lead ASO. Finally, we performed in vitro experiments to explore alternative splicing as a potential mechanism whereby HNRNPH2 controls the expression of HNRNPH1. We found that this mechanism is likely involved in upregulation of HNRNPH1 in human WT iPSC-derived neurons upon knockdown of *HNRNPH2* by a human *HNRNPH2* research ASO. For all in vivo studies, experimental sample sizes were based on previously published phenotyping data and the number of animals per treatment group is included in the figure legend. Data collection was stopped once predetermined *Hnrnph2* mutant group sizes were reached. For ddRT-PCR experiments and western blots, 3-4 biological repeats were run. For rescue studies, litters of mice were randomly assigned to treatment groups and dosed with the same compound (control ASO or *Hnrnph2* ASO). Researchers were not blinded to the genotype or treatment condition in the in vivo experiments. Litters containing no *Hnrnph2* mutant mice (WT only litters) were excluded from the study. For the elevated plus maze, mice that fell from the maze were excluded from the data analysis of that test. In the juvenile treatment rescue study, all mice that experienced post-ICV seizures were excluded from subsequent behavioral analysis to minimize potential confounding effects of acute ASO toxicity on the phenotypes being evaluated. No outliers were removed from the data.

### Statistics

For ASO potency and duration of action on *Hnrnph2* and *Hnrnph1* mRNA levels in WT C57BL/6J mice, we used two-way ANOVA (dose/age harvested, treatment, dose/age harvested x treatment interaction) followed by Tukey’s test to compare between control ASO and *Hnrnph2* ASO at each dose/age harvested, between lowest control ASO dose and higher control ASO doses, and between lowest *Hnrnph2* ASO dose and higher *Hnrnph2* ASO doses. For experiments comparing the effect of control ASO, *Hnrnph2* ASO, and vehicle on mRNA or protein expression, we used a one-way ANOVA followed by Tukey’s multiple comparison test. For body weight, we used a used two-way ANOVA (sex, treatment, sex x treatment interaction) followed by Tukey’s test to compare males vs. females for each treatment, and control ASO vs. *Hnrnph2* ASO, control ASO vs. vehicle, and *Hnrnph2* ASO vs. vehicle in males and females separately.

The log-rank (Mantel-Cox) test was used to determine significant differences between Kaplan-Meier survival curves of all groups. If the test showed a significant difference, pairwise survival curve comparison was performed for control ASO-treated WT vs. control ASO-treated mutant, *Hnrnph2* ASO-treated WT vs *Hnrnph2* ASO-treated mutant, WT control ASO-treated vs. WT *Hnrnph2* ASO-treated, mutant control ASO-treated vs. mutant *Hnrnph2* ASO-treated mice and resulting *P* values were adjusted for multiple comparison by the Holm-Šídák method.

For experiments comparing control ASO vs. *Hnrnph2* ASO-treated mutants and their WT littermates where a single measurement was taken, we used two-way ANOVA (genotype, treatment, genotype x treatment interaction) followed by Tukey’s test or Sidak’s test to compare control ASO-treated WT vs. control ASO-treated mutant, *Hnrnph2* ASO-treated WT vs. *Hnrnph2* ASO-treated mutant, WT control ASO-treated vs. WT *Hnrnph2* ASO-treated, mutant control ASO-treated vs. mutant *Hnrnph2* ASO-treated mice. Only seizure score was analyzed by non-parametric Scheirer–Ray–Hare test, followed by Mood’s median test was used to perform group wise comparisons. For experiments comparing control ASO vs *Hnrnph2* ASO-treated mutants and their WT littermates where multiple measurements were taken over time, we used a linear mixed effects model with random intercept to explore how the measurement changed over time and how it was affected by genotype and treatment. The post hoc comparisons were adjusted by the false discovery rate method within each time period.

For the incidence of hydrocephalus and post ICV seizures, we used a 4-by-2 way and 4-by-3 way contingency table, respectively, followed by Fisher’s exact test. If the *P* value was significant, we performed pairwise comparisons of control ASO-treated WT vs. control ASO-treated mutant, *Hnrnph2* ASO-treated WT vs. *Hnrnph2* ASO-treated mutant, WT control ASO-treated vs. WT *Hnrnph2* ASO-treated, mutant control ASO-treated vs. mutant *Hnrnph2* ASO-treated groups, followed by Fisher’s exact test. *P* values were adjusted for multiple comparison by the Holm-Šídák method.

For experiments in HEK293T cells and iPSC-derived neurons, we used student t-test, or ordinary one-way ANOVA with Dunnett’s multiple comparison.

## List of Supplementary Materials

Materials and Methods

Fig. S1 to S9

Tables S1 to S7

References

## Acknowledgements

We thank members of the St. Jude Center for In Vivo Imaging and Therapeutics, including Melissa Johnson, Katie Ray, and Thomas Confer for ICV injection of ASOs. We thank the St. Jude Center for Advanced Genome Engineering, including Baranda Hansen and Shondra Pruett-Miller, for assistance with CRISPR/Cas9-modified cell lines. We thank Natalia Nedelsky for editorial assistance.

## Author contributions

For design and validation of the murine *Hnrnph2* and human *HNRNPH2* ASOs, JO and BP designed, performed, and analyzed laboratory experiments. BP contributed to drafting the manuscript. BP and CFB contributed to supervision of the overall study. For preclinical studies using these ASOs, AK, XY, OO, AS, and AJL designed and performed laboratory experiments. AK, YDW, AMK, TP, HJK, and JPT analyzed data. AK, HJK, and JPT drafted and revised the manuscript. HJK and JPT supervised the overall study.

## Data and materials availability

All data are available in the main text or as supplemental data.

## Supplementary Materials for

### Materials and Methods

#### Antisense oligonucleotide design and screening

Synthesis and purification of all chemically modified oligonucleotides was performed as previously described (*1*, *2*). The 2′-*O*-methoxyethylribose (MOE) gapmer ASOs were 20 nucleotides long, with a central gap segment comprising ten 2′-deoxynucleotides flanked on the 5′ and 3′ ends by five 2′-MOE modified nucleotides. All internucleoside linkages were phosphorothioate linkages, except those shown as ‘o’ in the sequences below, which are phosphodiester. Lyophilized ASOs were formulated in PBS without Ca^2+^ and Mg^2+^ (Gibco, 14190).

As previously described (*2*), ASOs targeting murine *Hnrnph2* or human *HNRNPH2* were designed at Ionis to the murine and human *hnRNPH2* pre-mRNAs, targeting both exonic and intronic sequences. More than 450 ASOs were designed to the murine *Hnrnph2* pre-mRNA, which were screened in primary mouse embryonic cortical cultures isolated from 16-day pups by adding the ASOs to the culture medium (free uptake). A similar number of ASOs were designed to the human *HNRNPH2* pre-mRNA and screened by free uptake in A431 cells (ATTC). The level of target mRNA reduction was assessed by RT-qPCR and normalized to total RNA level (RiboGreen, Quant-iT). Approximately 40 leads from both the human and mouse screens were evaluated in a 4 point dose-response study to confirm activity and to rank the leads. The top 19 murine *Hnrnph2* ASOs that showed the most efficient knockdown of *Hnrnph2* expression compared to untreated controls were then injected intracerebroventricularly in an 8-week, 700 µg activity and tolerability study in 8–12-week-old female C57BL/6J mice. A dose of 700 µg was selected based on prior work showing that this dose generally achieves greater than 70% inhibition for active ASOs and will identify poorly tolerated ASOs. PBS was used as vehicle control. Mice were observed acutely for any adverse effects as previously described (*3*, *4*) and subsequently weighed and assessed for overall health weekly for 8 weeks. At the end of 8 weeks, animals were sacrificed, tissues collected, and RNA extracted to support analysis of ASO activity and tolerability, measuring astrocyte and microglial activation markers. Screening data for 3 lead murine *Hnrnph2* ASOs is shown in table S1. Similar screens were performed for the ASOs targeting human *HNRNPH2*. Levels of *Hnrnph1, Hnrnph2*, and *Aif1* were assessed by RT-qPCR. TaqMan primer/probe sets were employed with 56-FAM_Zen_3IABkFQ probe chemistry (Integrated DNA Technologies). Forward (F), reverse (R), and probe (P) sequences are as follows. Murine *Hnrnph1 –* F: CCTTCGTGCAGTTTGCTTC, R: GATCATAATGAGTTCTAACTTCAGCTC, P: ACCTGTGCCCTATTCTTTCCTTGTGTT; Murine *Hnrnph2* is captured in table S4; Murine *Gapdh* – F: GGCAAATTCAACGGCACAGT, R: GGGTCTCGCTCCTGGAAGAT, P: AAGGCCGAGAATGGGAAGCTTGTCATC.

The sequences of the selected ASOs are as follows: mouse non-targeted control ASO, 5′-CCTATAGGACTATCCAGGAA-3′; mouse *Hnrnph2* ASO1, 5′-GCTTATACCATTTCTCTTAT-3′. Human non-targeted control ASO, 5′-AATGCTTTCATAACTTCCAC-3′; human *HNRNPH2* ASO2, 5′-GTACAGCTTTTATATCTGGA-3′.

#### Animals

WT C57BL/6J mice were used for ASO dose response and longitudinal studies, distribution, and tolerance studies, with a balance of males and females. For all other studies, *Hnrnph2* P209L or R206W mutant knockin mice (*5*) were used, and male and female cohorts were analyzed separately. Heterozygous mutant females were bred to WT C57BL/6J males to generate heterozygous mutant females (*Hnrnph2^P209L/X^*), hemizygous mutant males (*Hnrnph2R^206W/Y^*, *Hnrnph2^P209L/Y^*), and their WT female (*Hnrnph2^X/X^*) and male (*Hnrnph2^X/Y^*) littermates. All experiments were performed on generation F12 or later. Animals were group housed under standard conditions. Experimental protocols were approved by the Institutional Animal Care and Use Committee at St. Jude Children’s Research Hospital.

#### Intracerebral ventricular injection of ASOs

Litters were randomly assigned to treatment groups. Control or *Hnrnph2* ASO was diluted to the appropriate concentration in PBS with 0.0135% Fast Green to aid in verification of the injection location. For neonatal treatment, postnatal day 2 to 4 pups were cryo-anesthetized for 5-10 minutes. The scalp was cleaned with 70% ethanol and freehand injections were performed using a custom 10 µl Hamilton syringe with 32-gauge needle (80308, Hamilton). Using the bregma and lambda as anatomical references, 3 µl control ASO, *Hnrnph2* ASO, or vehicle was injected freehand into the left ventricle. After injection, pups were rewarmed in a petri dish with bedding from the dam’s cage until the entire litter was injected. When pups were warm, pink, and capable of spontaneous movement, the litter was sprinkled in dam urine, returned to the home cage, monitored to ensure maternal acceptance, and observed closely for 24 hours post procedure. For juvenile treatment, postnatal day 27 to 29 weanlings were anesthetized with isoflurane by inhalation and their hair shaved with clippers to expose the scalp. After the scalp was cleaned with 70% ethanol, lidocaine/bupivacaine was injected subcutaneously, and a sagittal incision was made to expose the skull. Using the bregma and lambda as anatomical references, 3 µl control ASO, *Hnrnph2* ASO, or vehicle (0.0135% Fast Green in PBS) was injected freehand into the left ventricle using a custom 10 µl Hamilton syringe with 32 gauge needle (80308, Hamilton). After injection, the incision was closed with tissue adhesive and mice were moved to a heating pad where they were monitored for recovery from anesthesia and acute post-surgery complications. Once fully awake, mice were returned to their home cage with littermates where they were received subcutaneous meloxicam injections every 24 hours for 3 days and monitored for post-operative complications twice daily for 1 week.

#### Differentiation and ASO treatment of WT human iPSC-derived neurons

Human male BJFF.6 iPSCs were differentiated into cortical neurons with a two-step protocol (pre-differentiation and maturation) as previously described (*6*). When iPSCs reached 70-80% confluence, cells were washed twice with DPBS and dissociated with Accutase (STEMCELL Technologies) and collected cells were filtered using a cell strainer (STEMCELL Technologies). Filtered cells were centrifuged at 200 rcf for 5 min at room temperature and pellets were resuspended with medium containing N2 supplement, non-essential amino acids (NEAA), GlutaMAX Supplement and Y-27632 (STEMCELL Technologies) and 1 μg/ml doxycycline hyclate (Sigma Aldrich). Cell counting was performed using 10 μl and a Countess cell counter (Thermo Fisher). Cells were subplated at 2 x 10^6^ cells/well in 6-well dishes coated with Geltrex (Gibco) in knockout Dulbecco’s modified Eagle’s medium (KO-DMEM)/F12. The medium was changed daily for 3 days and Y-27632 was removed from day 2. For maturation, pre-differentiated precursor cells were washed, dissociated, counted, and subplated at 25 x 10^4^ cells/ml on dishes coated with 50 μg/ml poly-L-ornithine and 1-2 μg/cm^2^ laminin (Millipore Sigma) in BrainPhys neuronal medium (STEMCELL Technologies) containing N2 (Thermo Fisher Scientific), B-27, 20 ng/ml BDNF (PeproTech), 20 ng/ml GDNF (PeproTech), 500 μg/ml dibutyryl cyclic-AMP (Sigma Aldrich), 200 nM L-ascorbic acid (Sigma Aldrich), 1 μg/ml natural mouse laminin (Thermo Fisher Scientific), 1 μM AraC (Sigma Aldrich), Y-27632 (STEMCELL Technologies) and 1 μg/ml doxycycline hyclate. Y-27632 was removed from day 2. Half-medium was changed every other day. iPSC-derived neurons were treated with *HNRNPH2* ASO or vehicle (DPBS) from day 7 to day 21 of neuronal maturation. The ASO was initially added directly to the media to a final concentration of 10 µM, then at each subsequent half-media change to achieve the same final concentration. RNA isolation, protein isolation, and Western blotting in iPSC-derived neurons were performed as described for HEK293T cells below. ddRT-PCR was performed as described for mouse tissue below.

#### Cell culture, transfection, Western blot, and digital droplet RT-PCR

HEK293T cells were maintained in Dulbecco’s Modified Eagle’s Medium (DMEM) supplemented with 10% fetal bovine serum. Cells were transfected with RNAiMAX reagent (Thermo Fisher Scientific, 13778075) according to the manufacturer’s protocol, using either ON-TARGETplus Non-targeting Pool siRNA (Dharmacon, D-001810-10-05) or ON-TARGETplus human hnRNPH2 siRNA (Dharmacon, L-013245-02-0005). For plasmid-based transfections, cells were transfected with HA-minigene and/or GFP-hnRNPH2 using Lipofectamine 3000 (Thermo Fisher Scientific, L3000015) according to the manufacturer’s instructions. Twenty-four hours post-transfection, cells were lysed on ice for 30 min in lysis buffer (25 mM Tris-HCl pH 7.5, 150 mM NaCl, 1 mM EDTA, 1% NP-40, 5% glycerol [v/v]) supplemented with 1× Complete Protease Inhibitor Cocktail (Thermo Fisher Scientific, 1862209). An aliquot to 3% of the lysate was used as input. Proteins were resolved by electrophoresis on NuPAGE Novex 4–12% Bis-Tris gels (Invitrogen) and transferred to membranes. Membranes were blocked with Intercept TBS blocking buffer (LI-COR, 927-60001) and incubated with primary antibodies followed by species-specific IRDye secondary antibodies. Imaging was performed on an Odyssey CLx system and analyzed on Image Studio software version 5.5 (LI-COR). Details on antibodies used in western blots are provided in table S5. For RNA isolation, cells were washed twice with DPBS and processed following the manufacturer’s instructions using the RNeasy Mini Kit (74106, Qiagen). RNA was quantified using the Quant-iT RiboGreen RNA Assay Kit (R11490, Thermo Fisher Scientific), or NanoDrop 8000 Spectrophotometer (Thermo Fisher Scientific). ddRT-PCR was performed as described for mouse tissues below. Target and reference genes with assay details are listed in table S4.

#### RT-PCR, digital droplet RT-PCR, and Western blot in mouse tissue

Mice were anesthetized by isoflurane inhalation and whole brains were harvested and cut in half sagitally before being flash frozen in liquid nitrogen and stored at −80°C until use. For RNA isolation from mouse brain, tissue was treated with RNAlater-ICE (AM7030, Thermo Fisher Scientific) to prevent degradation, processed using the RNeasy Midi Kit with on-column DNase digestion (75144, Qiagen), and quantified with the Quant-it RiboGreen RNA Assay Kit (R11490, Thermo Fisher Scientific). For gene expression by RT-PCR, 400 ng of RNA from mouse brain was converted to cDNA with the SuperScript IV First-Strand Synthesis System (18091050, Thermo Fisher Scientific) using random hexamer primers. Two µl of cDNA was used for PCR with the Platinum II Hot-Start PCR Master Mix (14001012, Thermo Fisher Scientific). Products were resolved on a 3 % agarose gel, imaged on the ChemiDoc MP system (Bio-Rad), and densitometry per formed with ImageJ. For gene expression analysis by digital droplet PCR, 1-10 ng RNA (depending on the target transcript) was used with a one-step RT-ddPCR Advanced Kit for Probes (1864021, Bio-Rad). Data were analyzed with the QX Manager Software Version 2.0 (Bio-Rad), copies were normalized to those of a reference gene with similar expression level, and expression plotted as a percentage of control. Target and reference genes with assay details are listed in table S4.

For western blots in mouse, brain tissue was processed for total cell RIPA-soluble fractions as previously published (*7*), or for cytoplasmic and nuclear fractions with the Subcellular Protein Fractionation Kit for Tissues (87790, Thermo Fisher Scientific) according to the manufacturer’s protocol. Protein concentrations were determined by BCA protein assay (A55864, Thermo Fisher Scientific), and 15 μg total cell RIPA-soluble fraction, 100 μg cytoplasmic fraction, or 85 μg nuclear fraction was loaded onto 8% WedgeWell gels (NW00080BOX, Thermo Fisher Scientific). Electrophoresis was performed using the Bolt Bis-Tris mini gel system (Thermo Fisher Scientific) and proteins were transferred to PVDF membranes (IB24001, Thermo Fisher Scientific) using the iBlot 2 dry blotting system (Thermo Fisher Scientific). Membranes were blocked with Intercept TBS blocking buffer (LI-COR, 927-60001) and incubated with primary antibodies followed by species-specific IRDye secondary antibodies. Imaging was performed on an Odyssey CLx system and analyzed on Image Studio software version 5.5 (LI-COR). Details on antibodies used in western blots are provided in table S5.

#### RNA-seq data analysis

Total RNA was extracted with an RNeasy Universal Midi kit (Qiagen; 75144) as described above. A sequencing library was prepared with a TruSeq Stranded Total RNA Kit (Illumina) and sequenced with an Illumina HiSeq system with 100-bp read length. Total stranded RNA sequencing data were processed by the internal AutoMapper pipeline. Raw reads were first trimmed (Trim-Galore v0.60), mapped to mouse (GRCm39) genome assembly (STAR v2.7) and then the gene level values were quantified (RSEM v1.31) based on GENCODE annotation (vM22). We obtained the TPM (transcript per million) counts for genes with TPM greater than one in at least one sample and the gene expression analysis was performed with non-parametric ANOVA using Kruskal-Wallis and Dunn’s tests on log-transformed TPM counts between three replicates of each experimental group, implemented in Partek Genomics Suite v7.0 software (Partek). The expression of a gene was considered significantly different if the adjusted (through two-stage linear step-up procedure of Benjamini, Krieger and Yekutieli) (*8*) *P* value < 0.05 (equivalent to FDR of 5%) and log2FC > 0.322. For alternative splicing analysis, the aligned and sorted BAM files after STAR alignment were used for AS analysis using rMATS (v4.0.2) (50). A3SS, A5SS, SE, RI, and MXE events were evaluated. Significant AS events from the two-group differential alternative splicing analysis were identified using the following criteria: average RNA-seq read count ≥10 in both sample groups; filter out events with average PSI value <0.05 or >0.95 in both sample groups; FDR ≤0.05; |ΔPSI| ≥0.05 (*9*).

#### Immunofluorescence

Mice were anesthetized by isoflurane inhalation and whole brains harvested, cut in half sagitally, postfixed in 10% neutral buffered formalin (NB0345, StatLab), and embedded in paraffin. Alternatively, mice were anesthetized by isoflurane inhalation and transcardially perfused with 4% paraformaldehyde (J19943, Thermo Fisher Scientific) before whole brains were harvested, cut in half sagitally, cryoprotected in 30% sucrose in PBS, and embedded in Tissue Freezing Medium (TFM-C, GeneralData). To assess the distribution and activity of the *Hnrnph2* ASO in the mouse brain, 4-µm sagittal paraffin sections were cut on a microtome, deparaffinized and subjected to proteinase K antigen retrieval (S3020, Dako) or heat-induced epitope retrieval (50-187-82, Thermo Fisher Scientific) depending on the primary antibodies used. Sections were then stained according to standard immunofluorescence protocols. For sections stained with both the rabbit anti-ASO and rabbit anti-HNRNPH2 antibodies, the anti-HNRNPH2 antibody was labeled with CF568 using a Mix-n-Stain antibody labeling kit (92235, Biotium). Details on antibodies used are provided in table S7.

For ASO distribution across the mouse brain, images were acquired with a Leica SP8 confocal microscope or a Lionheart slide scanner. For ASO distribution into brain cell types, images were acquired with a Leica Stellaris confocal microscope. For *Hnrnph2* downregulation in brain cells containing *Hntnph2* ASO, images were acquired with a Leica Stellaris confocal microscope or a Leica SP8 confocal microscope.

#### Histopathology

For histopathological evaluation of *Hnrnph2* ASO toxicity, mice were anesthetized by isoflurane, euthanized, perfused transcardially with 10% neutral buffered formalin and subjected to a standard full necropsy. Tissues were dissected, fixed in 10% neutral buffered formalin, and processed using the HistoCore PEGASUS Tissue Processor (Leica Biosystems). Processed tissues were embedded in paraffin and sectioned at a thickness of 4 µm using a HistoCore AUTOCUT fully automated rotary microtome (Leica Biosystems). Tissue sections were stained with hematoxylin and eosin (H&E) using the HistoCore SPECTRA ST Stainer (Leica Biosystems). Simplex immunohistochemistry was performed on sequential brain sections to detect CD68 (macrophages), CD163 (peripheral monocyte-derived macrophages), IBA1/AIF1 (microglia/macrophages), and GFAP (astrocytes). Details on epitope retrieval, primary antibodies, staining platform and detection system used are summarized in table S6. Sections were reviewed by a board-certified veterinary pathologist (AMK). Histopathologic lesions and positive reactivity for each immunohistochemistry assay were semi-quantitatively scored for severity (*10*). For GFAP and IBA1, positive reactivity above baseline expression in the brain was scored.

#### Audiogenic seizure susceptibility

Susceptibility to audiogenic seizures was tested as previously described (*5*). Briefly, mice were exposed to a 120-125 dB fire bell inside a clear acrylic box (30 x 30 x 30 cm) for 60 seconds and the intensity of the response (seizure severity score) was categorized as 0 for no response or slight startle, 1 for wild running, 2 for clonic seizures, 3 for tonic seizures, and 4 for respiratory arrest (*11*).

#### Motor function tests

Mice were tested for balance and coordination, muscle strength and gait performance as previously described (*5*). Rotarod analysis was performed on an accelerating rotarod apparatus (IITC Life Science) using a 2-day protocol consisting of a training day with rotation speed set at 4 rpm for 5 minutes, followed by a testing day with rotation speed set to accelerate from 4 to 40 rpm at 0.1 rpm/s. On test day, latency to fall was recorded for four separate trials per mouse, with a 15-minute rest period between each trial, and the average latency to fall across the 4 trials was calculated. The beam walking test was performed using a 2-day, multibeam (MazeEngineers) protocol (*12*). Total latency to cross all 3 beams and number of hind paw slips on the 12-mm square beam was analyzed. Grip strength was measured using a grip strength meter (Bioseb) as grams of force for front paws in six repeated measurements. The average grip strength across the 6 trials was calculated. For gait analysis, the front and hind paws of each animal were dipped in water-soluble non-toxic red and blue paint, respectively. The animal was then placed in a 70-cm long tunnel lined on the bottom with Whatman filter paper and allowed to walk through one time. Footprints were scanned and analyzed with Image J for stride length, fore- and hind base width, and overlap (*13*).

#### Cognitive function tests

Mice were assessed for anxiety in the elevated plus maze, anxiety and locomotion in the open field test, and spatial learning and memory in the Morris water maze using the ANY-maze automated activity monitoring system and software (v7.2) as previously described (*5*). Briefly, in the elevated plus maze (Stoelting), mice were placed in the center facing an open arm away from the investigator and allowed to explore the maze undisturbed for 5 minutes (*14*). Parameters including total distance traveled, time spent in open arms and time spent in the maze center were recorded and analyzed. The animal’s entire area was used to score arm entries and exits, with at least 80% of the animal needed for an arm entry to occur, and mice were considered to be in the center zone if not in any arm. Mice that fell off the maze were excluded from the analysis. For the open field test, mice were placed in the center of a 40 cm x 40 cm open field arena (Stoelting) facing away from the investigator and allowed to explore undisturbed for 20 minutes (*15*). Parameters including total distance traveled and time in the center zone (defined as the center 24 cm x 24 cm of the maze) were recorded and analyzed. The animal’s center was used to score zone entries and exits. The Morris water maze (MazeEngineers) consisted of a cued phase to test the physical ability of the mice to complete the task, followed by a training/spatial learning phase and finally, a spatial memory test phase (*16*). Parameters including mean speed, time in the thigmotaxis zone (within 10 cm of the pool wall), latency to find the visible platform (cued trials), latency to find the hidden platform (training trials), cumulative distance from the platform (training trials), latency to enter the platform location (test trial), and platform location crosses (test trial) were recorded and analyzed.

## Supplementary Figures

**Fig. S1.**
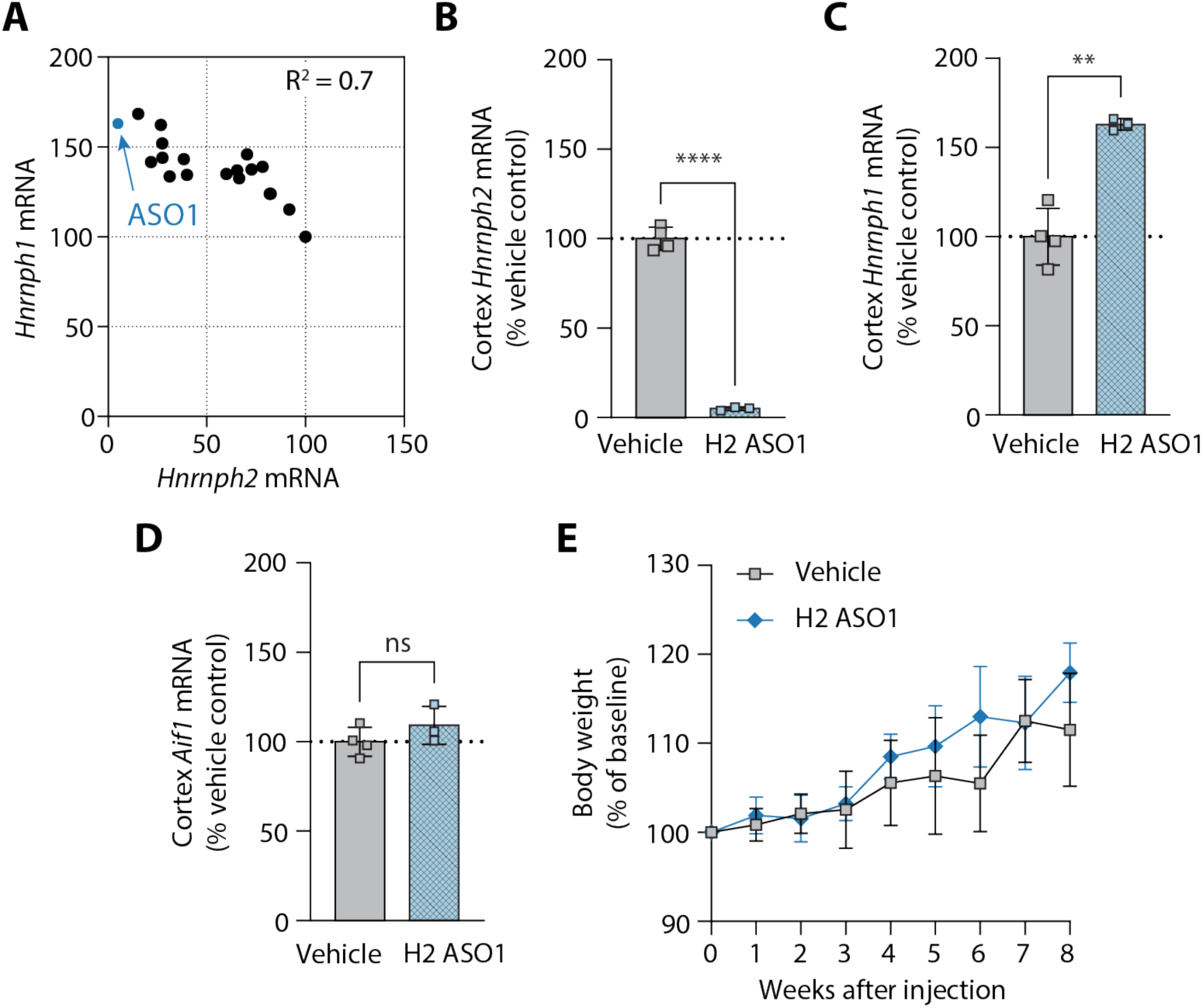
Initial characterization of mouse *Hnrnph2* ASOs in adult female C57BL/6J mice. (**A**) Expression levels of *Hnrnph2* and *Hnrnph1* in mouse cortex 8 weeks after an ICV injection with 700 µg of the tested *Hnrnph2*-targeted ASOs. Each dot represents a different ASO, with the blue dot indicating ASO1, which was used in all subsequent experiments. mRNA levels were measured by qRT-PCR, normalized to a reference gene, and expressed as a percentage of the vehicle (PBS) control level. There was a significant inverse relationship between *Hnrnph2* and *Hnrnph1* levels (*r* = −.8173, *P* < 0.0001, with an R^2^ value of 0.7, *n* = 3-4 mice per group). (**B-D**) Mouse *Hnrnph2* ASO1 knocks down *Hnrnph2* (B), increases *Hnrnph1* (C), and does not alter expression of microglial activation marker *Aif1* (D) in cortex 8 weeks following a single 700 µg ICV injection in adult WT mice. mRNA levels were measured by qRT-PCR, normalized to a reference gene, and expressed as a percentage of the vehicle (PBS) control level. \*\*\*\**P* < 0.0001, \*\**P* < 0.01, unpaired two-tailed t-test, *n* = 3-4 mice per group. Data are represented as mean ± SD. (**E**) Body weight was measured weekly across the study. No significant differences were observed in the body weight as a percentage of baseline between the vehicle- and *Hnrnph2* ASO1-treated mice across the study using multiple unpaired t-tests with two-stage step-up (*8*) multiple comparisons false discovery rate method.

**Fig. S2.**
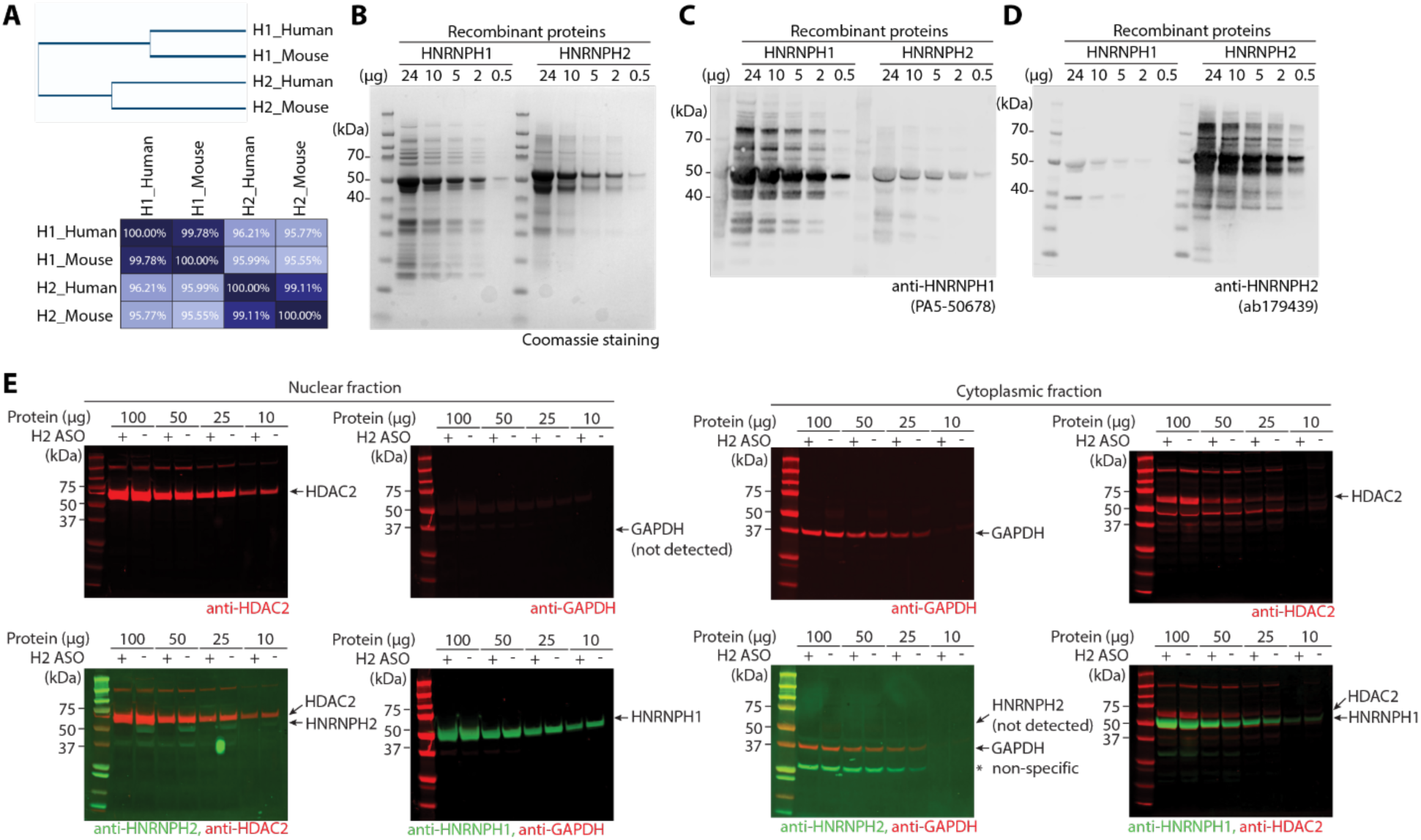
Validation of antibody specificity and efficiency of subcellular fractionation. (**A**) Phylogenetic tree of human and mouse HNRNPH1 and HNRNPH2 proteins. The table displays the percentage identity of their amino acid sequences. (**B-D**) Purified recombinant mouse HNRNPH1 and HNRNPH2 proteins were loaded onto gels in the indicated amounts. The gels were stained with Coomassie blue (B) or blots probed with the indicated antibodies (C, D). (**E**) Nuclear or cytoplasmic fractions prepared from mouse whole brains were loaded onto gels in the indicated amounts. Blots were probed with antibodies against HNRNPH2, a nuclear protein (HDAC2), and a cytoplasmic protein (GAPDH). Nuclear fractions showed a strong HDAC2 signal and no signal for GAPDH, supporting completeness of the fractionation. HNRNPH2 was detected in the nuclear fraction, and the level was reduced in *Hnrnph2* ASO-treated samples. HNRNPH1 was also detected in nuclear fractions. Cytoplasmic fractions showed a strong GAPDH signal as well as some signal for HDAC2, particularly when larger amounts of protein were loaded.

**Fig. S3.**
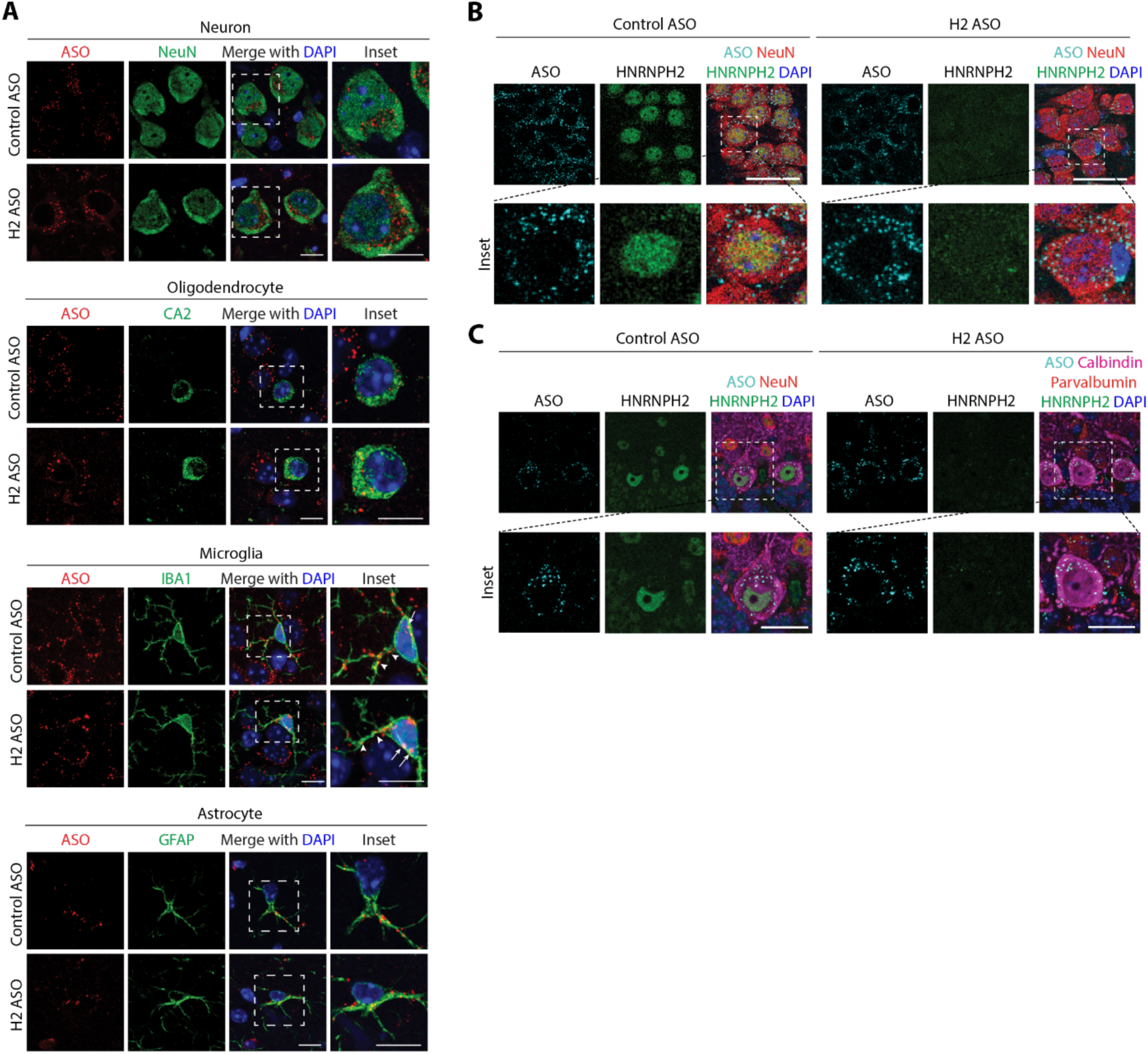
Distribution and activity of *Hnrnph2* ASO in the mouse brain. WT mice were treated with 30 µg control or *Hnrnph2* ASO in a single bolus ICV injection at PND 2-4. Brains were harvested at PND 21, processed for immunofluorescence, and slides imaged by confocal microscopy. (**A**) ASO signal was detected inside neocortical NeuN+ neurons, CA2+ oligodendrocytes, and IBA1+ microglia, as well as hippocampal GFAP+ astrocytes in both control ASO- and *Hnrnph2* ASO-treated samples. White arrows and arrowheads indicate ASO present in microglial cell bodies and processes, respectively. (**B**) HNRNPH2 signal was detected in nuclei of NeuN+ neurons in the hippocampus of control ASO-treated brains, but not in *Hnrnph2* ASO-treated brains. (**C**) HNRNPH2 signal was detected in nuclei of calbindin+ and parvalbumin+ Purkinje cells of the cerebellum in control ASO-treated brains, but not in *Hnrnph2* ASO-treated brains. Scale bars, 7 μm in (A) and (C), 20 μM in (B).

**Fig. S4.**
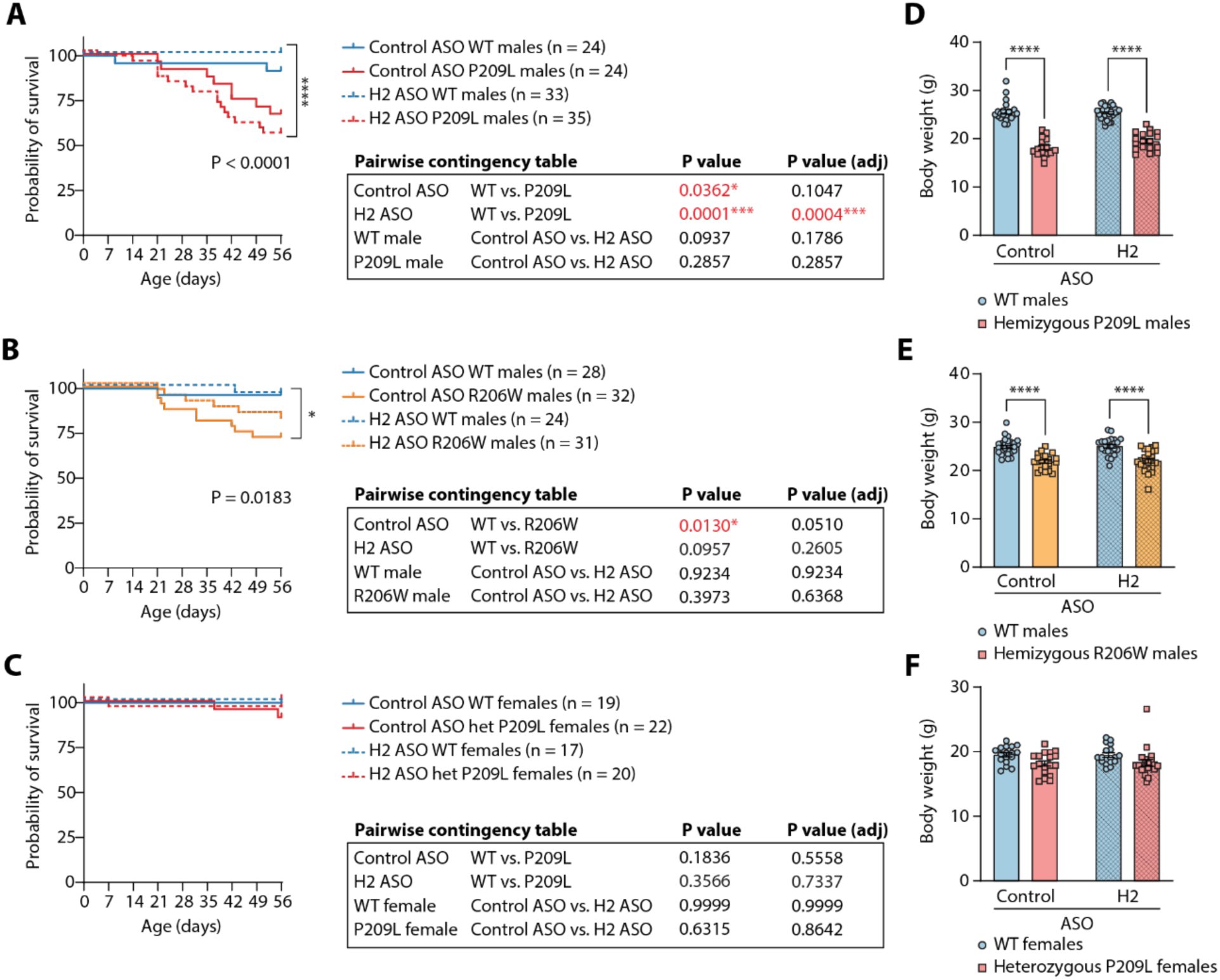
Neonatal treatment with the *Hnrnph2* ASO does not influence the survival and body weight phenotypes of mutant *Hnrnph2* mice. (**A-C**) Kaplan-Meier survival curves up to 8 weeks of age for P209L male mice (A), R206W male mice (B), and P209L female mice (C) treated with 30 µg control or *Hnrnph2* ASO by ICV injection at PND 2-4. \*\*\*\**P* < 0.0001, \**P* = 0.0183 by Mantel-Cox test. Table shows *P* values for pairwise survival curve comparisons after adjustment for multiple comparisons by the Holm-Sidak method. (**D-F**) Effect of neonatal ASO treatment on body weight of P209L male mice (D), R206W male mice (E), and P209L female mice (F) at 8 weeks of age. \*\*\*\**P* < 0.0001, 2-way ANOVA with Sidak’s multiple comparison. For (D): control ASO WT *n* = 22, control ASO P209L *n* = 16, *Hnrnph2* ASO WT *n* = 33, *Hnrnph2* ASO P209L *n* = 18. For (E): control ASO WT *n* = 27, control ASO R206W *n* = 21, *Hnrnph2* ASO WT *n* = 23, *Hnrnph2* ASO R206W *n* = 24. For (F): control ASO WT *n* = 17, control ASO P209L *n* = 18, *Hnrnph2* ASO WT *n* = 17, *Hnrnph2* ASO P209L *n* = 20. Data are represented as mean ± SEM.

**Fig. S5.**
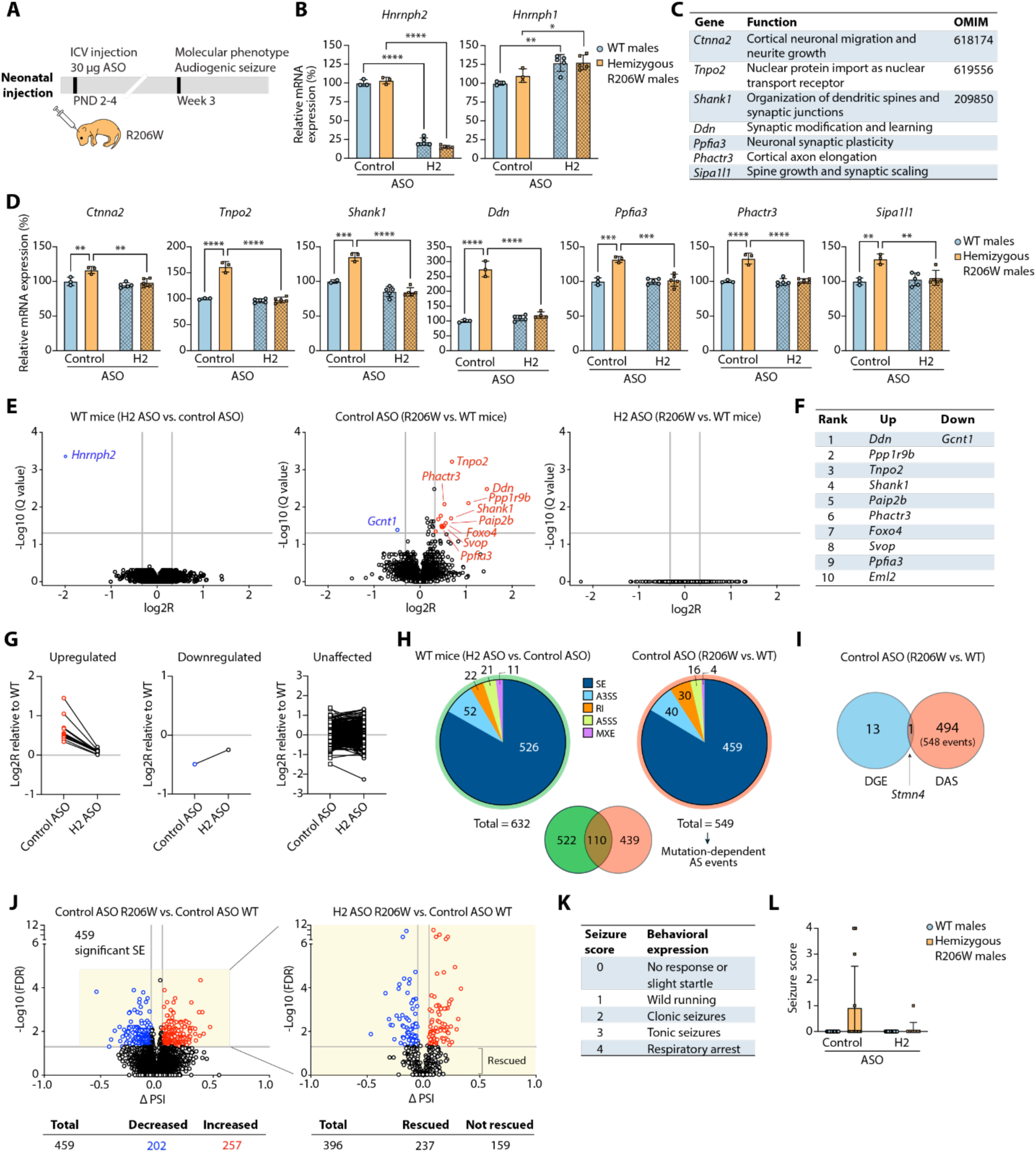
Neonatal treatment with *Hnrnph2* ASO rescues the molecular phenotype of juvenile R206W male mice. (**A**) Design of studies used to generate data in (B-L). All panels show data collected from R206W mutant mice or littermates at 3 weeks of age treated with 30 μg dose of ASO. (**B**) ddRT-PCR of whole brain tissues showing *Hnrnph2* and *Hnrnph1* mRNA levels. (**C**) Seven upregulated genes in both *Hnrnph2* mutant mice and human *HNRNPH2* mutant iPSC-derived neurons. (**D**) ddRT-PCR of whole brain tissues showing expression levels of genes listed in (C). (**E**) Volcano plots showing differentially expressed genes with significantly downregulated (blue) and upregulated (red) genes indicated. (**F**) Top 10 most upregulated genes and one downregulated gene by log2R from middle plot in (E). (**G**) Effects of *Hnrnph2* ASO on upregulated (left), downregulated (middle), and unchanged (right) genes at the individual gene level. (**H**) RNA-seq-derived differential alternative splicing events grouped by *Hnrnph2* ASO treatment (left) and R206W mutation (right). SE: skipped exon. A3SS: alternative 3′ splice site. RI: retained intron. A5SS: alternative 5′ splice site. MXE: mutually exclusive exon. Overlap between *Hnrnph2* ASO-dependent and R206W mutation-dependent events is shown. (**I**) Minimal overlap between R206W mutation-dependent events identified by differential gene expression (DGE) and differential alternative splicing (DAS) analyses. (**J**) Left, volcano plot showing the 459 significant skipped exon (SE) events in control ASO-treated R206W mutant mice with significantly increased (red) and decreased (blue) events. Right, volcano plot showing the effect of *Hnrnph2* ASO treatment on R206W mutation-dependent skipped exon events. (**K-L**) Audiogenic seizure susceptibility scoring (K) and assessment (L) of indicated mice. For panel B and D, 2-way ANOVA with Sidak’s multiple comparison, *n* = 3-4 mice per treatment. For panel L, non-parametric Scheirer–Ray–Hare test with Mood’s median test, control ASO WT males *n* = 11, control ASO R206W males *n* = 13, *Hnrnph2* ASO WT males *n* = 14, *Hnrnph2* ASO R206W males *n* = 13. For all statistical analyses, \*\*\*\**P* < 0.0001, \*\*\**P* < 0.001, \*\**P* < 0.01, \**P* < 0.05. Data shown as mean ± SEM.

**Fig. S6.**
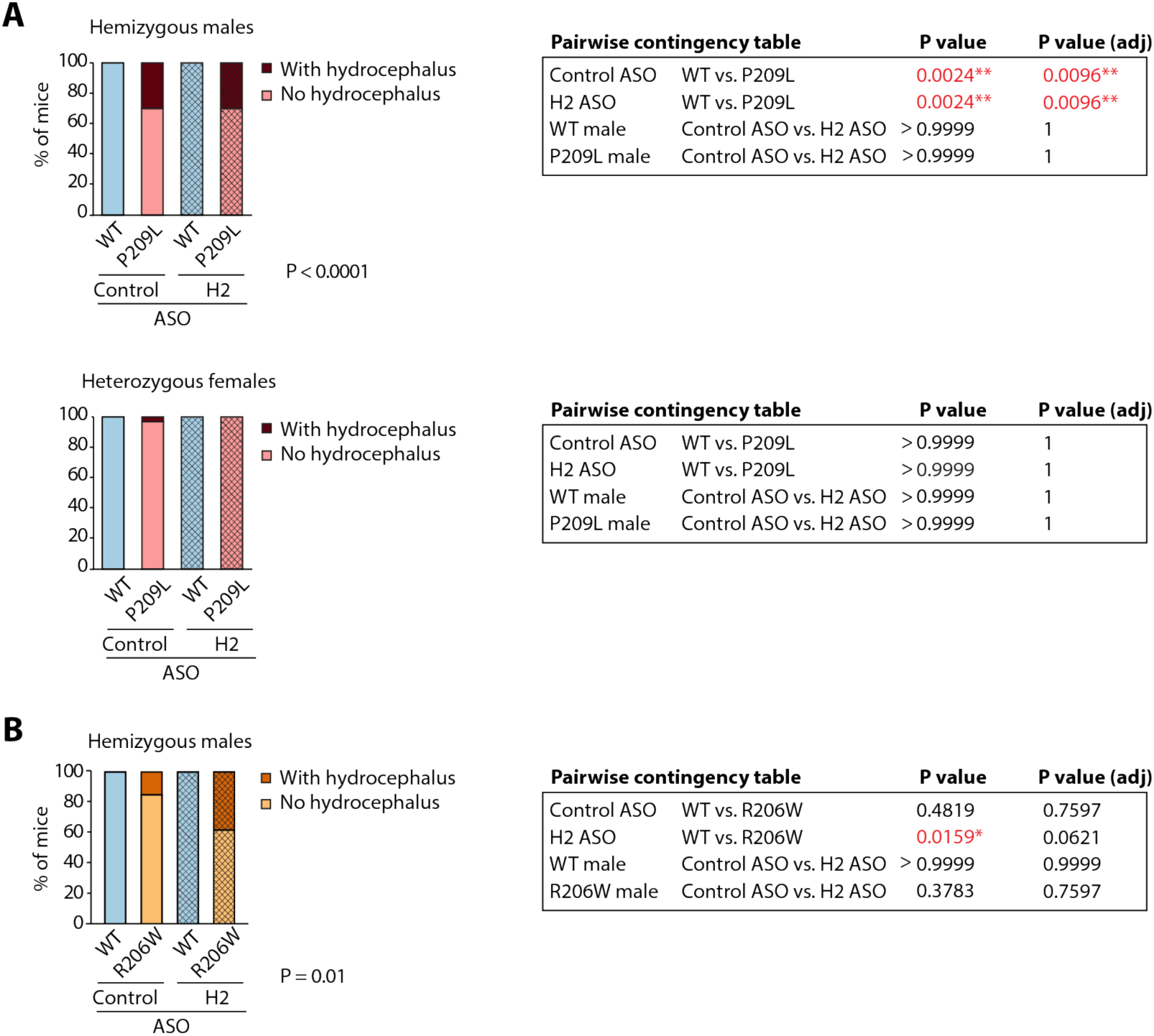
Effect of neonatal *Hnrnph2* ASO treatment on the incidence of hydrocephalus in male and female mice. Incidence of hydrocephalus observed at dissection of male and female P209L mice (**A**), and male R206W mice (**B**) at 3 weeks of age. *P* < 0.0001, *P* =0.01, Fisher’s exact test. For pairwise comparison, *P* values were adjusted for multiple comparisons by the Holm-Sidak method.

**Fig. S7.**
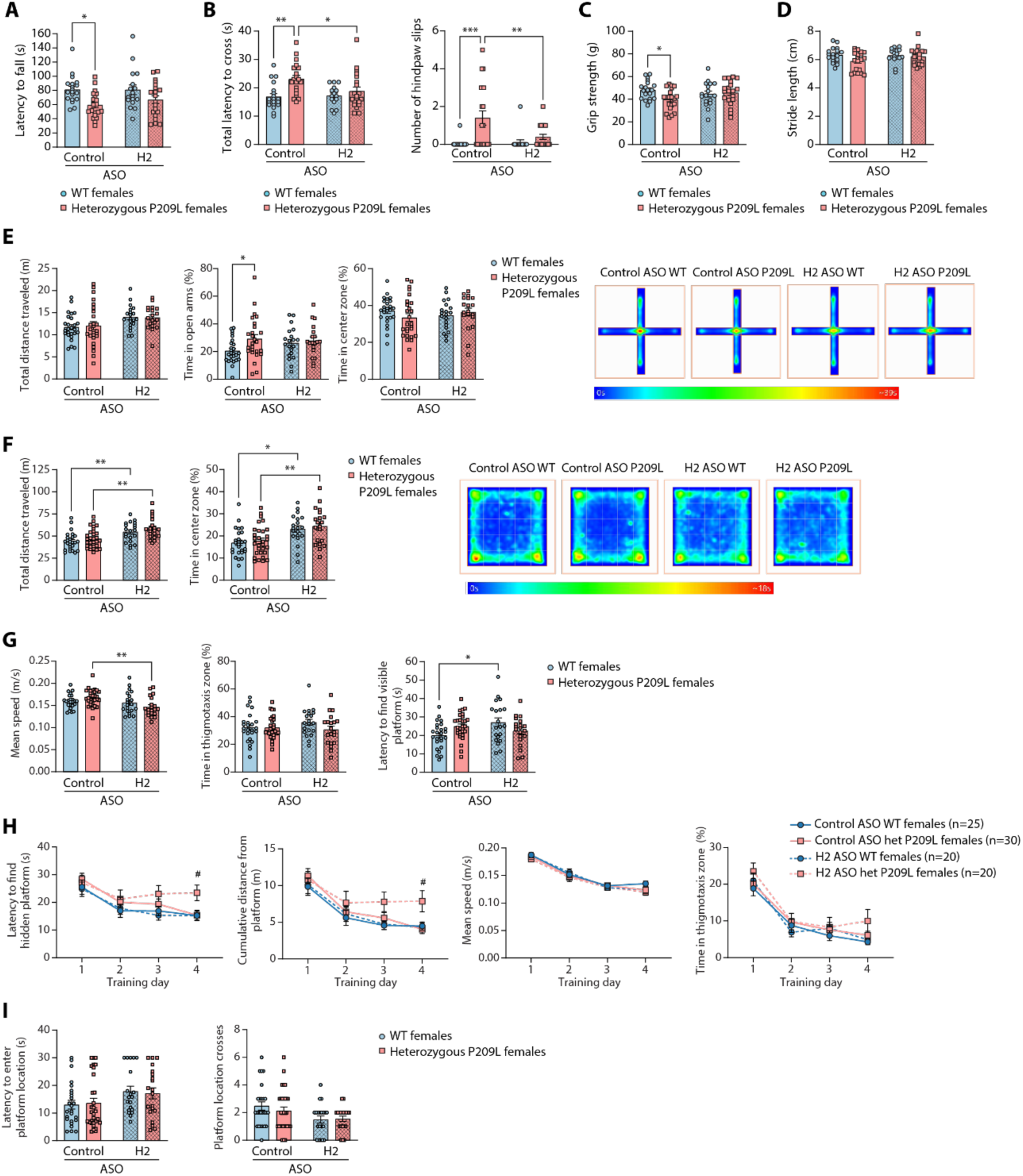
Impact of neonatal *Hnrnph2* ASO treatment on motor and cognitive functions in adult *Hnrnph2* P209L female mice. (**A-D**) Effect of neonatal *Hnrnph2* ASO treatment on rotarod performance (A), balance beam performance (B), grip strength (C), and stride length (D) in female P209L mice. (**E-I**) Effect of neonatal ASO treatment on elevated plus maze behavior (E), open field behavior (F), and Morris water maze performance (G-I) in female P209L mice. In panel E, heatmaps show the average animals’ center point for the groups for the 5-minute test. In panel F, heatmaps show the average animals’ center point for the groups for the 20-minute test. For panels A-G, \*\**P* < 0.01, \**P* < 0.05 by 2-way ANOVA with Sidak’s multiple comparison. For panel H, ^#^*P* < 0.05 by linear mixed effects model with random intercept. Post-hoc comparisons adjusted by FDR within each time point. For panels A-D, control ASO WT females *n* = 19, control ASO P209L females *n* = 20, *Hnrnph2* ASO WT females *n* = 17, *Hnrnph2* ASO P209L females *n* = 20. For panel B: control ASO WT females *n* = 19, control ASO P209L females *n* = 20, *Hnrnph2* ASO WT females *n* = 15-17, *Hnrnph2* ASO P209L females *n* = 20. For panel E: control ASO WT females *n* = 26, control ASO P209L females *n* = 26, *Hnrnph2* ASO WT females *n* = 20, *Hnrnph2* ASO P209L females *n* = 19. For panel F: control ASO WT females *n* = 26, control ASO P209L females *n* = 30, *Hnrnph2* ASO WT females *n* = 20, *Hnrnph2* ASO P209L females *n* = 20. For panel G-I: control ASO WT females *n* = 25, control ASO P209L females *n* = 30, *Hnrnph2* ASO WT females *n* = 20, *Hnrnph2* ASO P209L females *n* = 20. Data are represented as mean ± SEM.

**Fig. S8.**
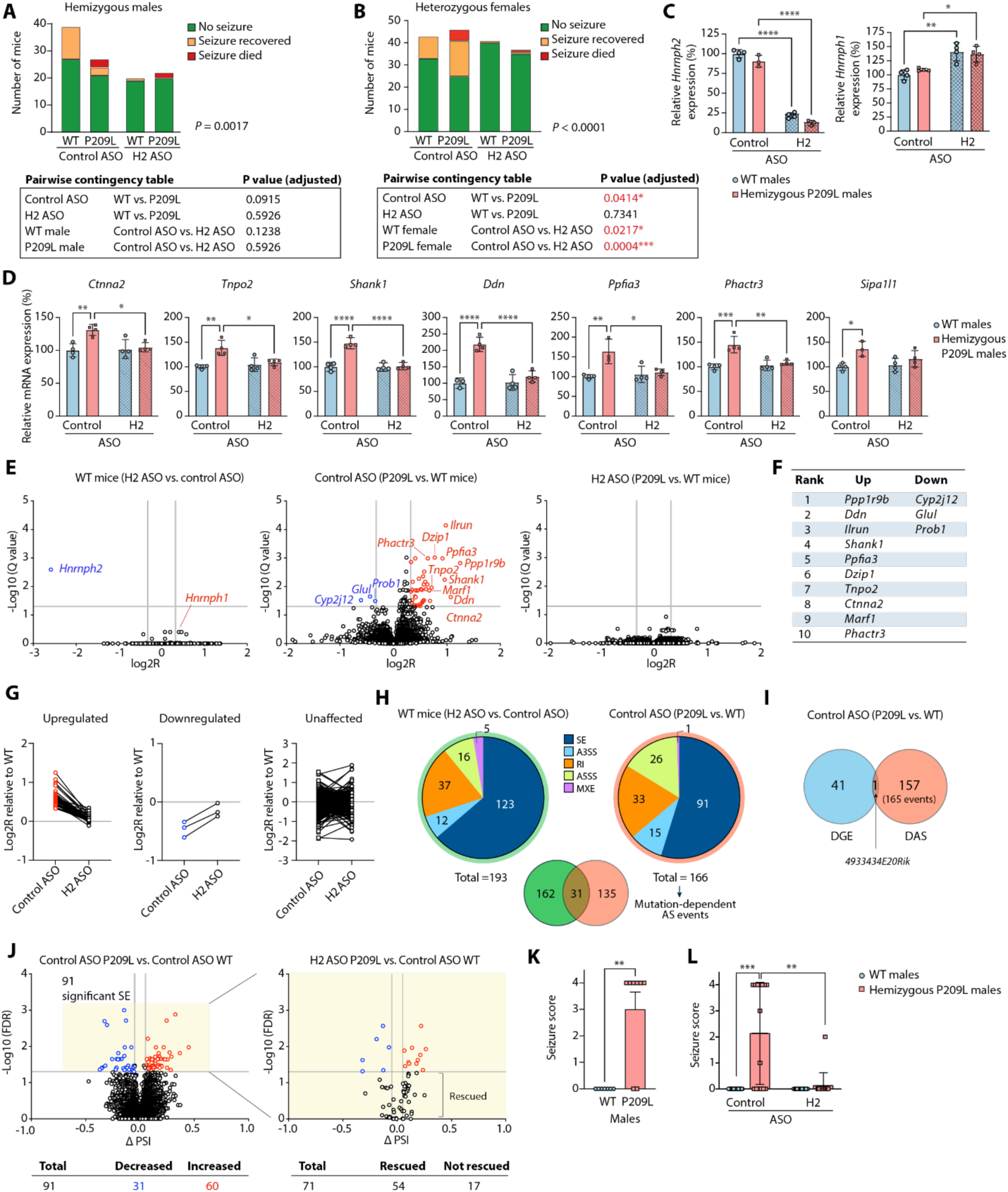
Juvenile ASO treatment is associated with post-ICV injection seizures and *Hnrnph2* ASO rescues the molecular phenotype of juvenile R206W male mice. (**A-B**) Effect of juvenile ASO treatment on the incidence of post-ICV injection seizures in male (A) and female (B) P209L mice. *P* values next to each graph were determined by Fisher’s exact test. Table shows *P* values for pairwise comparisons of contingency tables after adjustment for multiple comparisons by the Holm-Sidak method. For (A), control ASO WT males *n* = 39, control ASO P209L males *n* = 27, *Hnrnph2* ASO WT males *n* = 20, *Hnrnph2* ASO P209L males *n* = 22. For (B), control ASO WT females *n* = 43, control ASO P209L females *n* = 46, *Hnrnph2* ASO WT females *n* = 41, *Hnrnph2* ASO P209L females *n* = 37. (**C**) ddRT-PCR of whole brain tissues showing *Hnrnph2* and *Hnrnph1* mRNA levels. (**D**) ddRT-PCR of whole brain tissues showing expression levels of seven genes previously found to be upregulated in both *Hnrnph2* mutant mice and human *HNRNPH2* mutant iPSC-derived neurons. (**E**) Volcano plots showing differentially expressed genes with significantly downregulated (blue) and upregulated (red) genes indicated. (**F**) Top 10 most upregulated and 3 downregulated genes by log2R from middle plot in (E). (**G**) Effects of *Hnrnph2* ASO on upregulated (left), downregulated (middle), and unchanged (right) genes at the individual gene level. (**H**) RNA-seq-derived differential alternative splicing events grouped by *Hnrnph2* ASO treatment (left) and P209L mutation (right). SE: skipped exon. A3SS: alternative 3′ splice site. RI: retained intron. A5SS: alternative 5′ splice site. MXE: mutually exclusive exon. Overlap between *Hnrnph2* ASO-dependent and P209L mutation-dependent events is shown. (**I**) Minimal overlap between P209L mutation-dependent events identified by differential gene expression (DGE) and differential alternative splicing (DAS) analyses. (**J**) Left, volcano plot showing the 91 significant skipped exon (SE) events in control ASO-treated P209L mutant mice with significantly increased (red) and decreased (blue) events. Right, volcano plot showing the effect of *Hnrnph2* ASO treatment on P209L mutation-dependent skipped exon events. (**K-L**) Audiogenic seizure susceptibility in untreated (K) and ASO-treated (L) male mice. For panels C and D, 2-way ANOVA with Sidak’s multiple comparison, *n* = 3-4 mice per group. For panel K, non-parametric Mann-Whitney U test, WT males *n* = 7, P209L males *n* = 8. For panel L, non-parametric Scheirer–Ray–Hare test with Mood’s median test, control ASO WT males *n* = 25, control ASO P209L males *n* = 15, *Hnrnph2* ASO WT males *n* = 18, *Hnrnph2* ASO P209L males *n* = 16. For all statistical analyses, \*\*\*\**P* < 0.0001, \*\*\**P* < 0.001, \*\**P* < 0.01, \**P* < 0.05. Panels C-L show data collected from P209L mutant male mice or littermates at 8 weeks of age treated with 200 μg dose of ASO. Data shown as mean ± SEM.

**Fig. S9.**
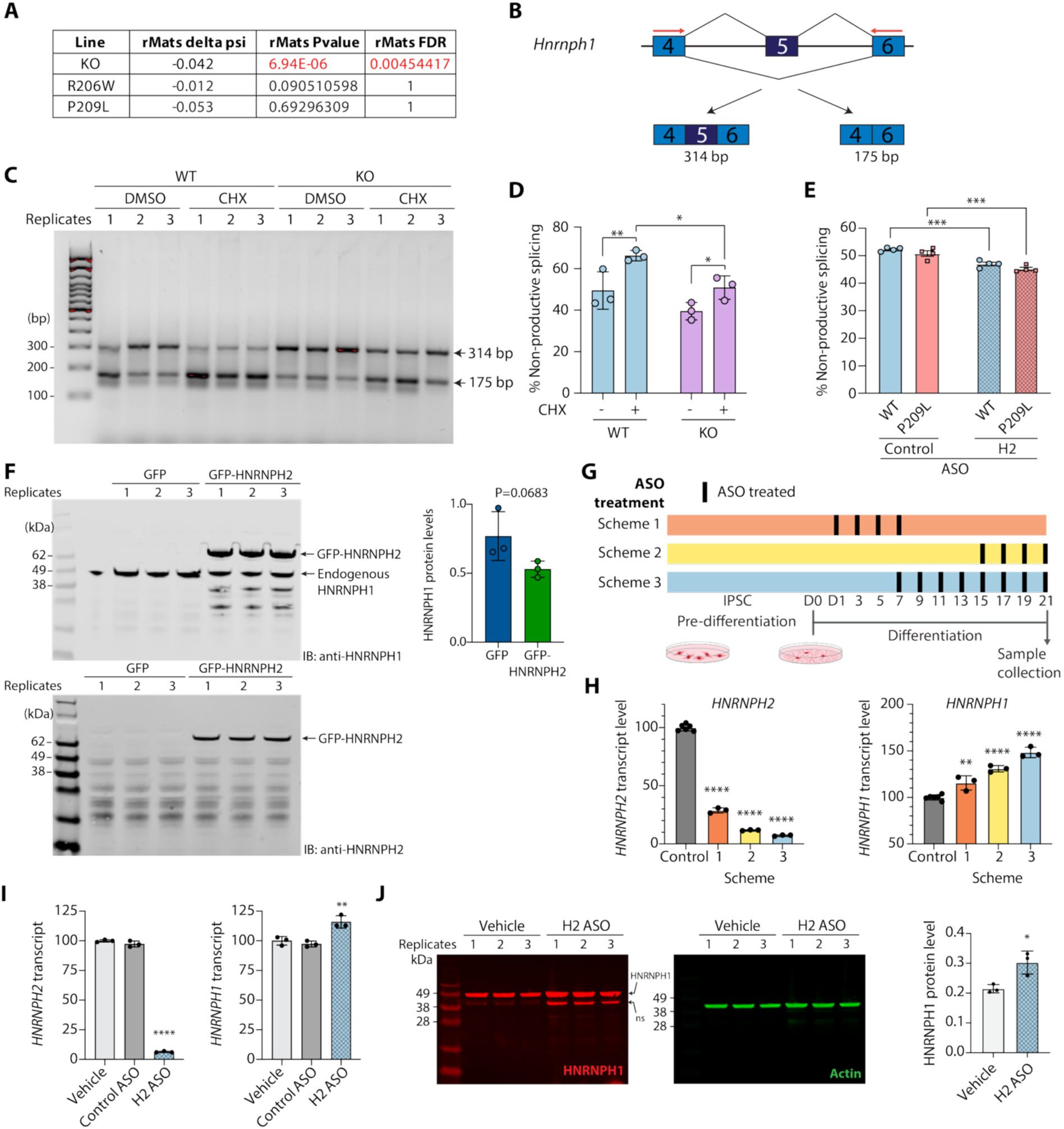
HNRNPH2 protein regulates the expression of *HNRNPH1* by modulating its splicing. (**A**) Significant skipping of essential exon (exon 5) of *Hnrnph1* in cortex from *Hnrnph2* KO mice, but not *Hnrnph2* P209L and R206W mutant mice, as detected by alternative splicing analysis of RNA-seq data (*5*). (**B**) Schematic representation of intron-exon structure of murine *Hnrnph1*, including the 139-nt essential exon and flanking exons. Forward and reverse primers were located in exon 4 and 6 (red arrows) and expected sizes of transcripts with and without the essential exon were 314 bp and 175 bp, respectively. (**C**) Alternative splicing of the essential exon of murine *Hnrnph1* was tested by RT-PCR in the absence (DMSO) or presence of cycloheximide (CHX) in mouse primary cortical neurons. Band identity was confirmed by sequencing. (**D**) Quantification of band intensity in (C) measured by densitometry. The percentage of non-productive splicing, calculated as the level of *Hnrnph1* transcripts lacking the essential exon divided by the level of total *Hnrnph1* transcripts (with and without the essential exon) was significantly increased after cycloheximide treatment in both WT and *Hnrnph2* KO neurons. After cycloheximide treatment, there was a significant reduction in non-productive splicing in *Hnrnph2* KO neurons compared to WT. (**E**) Alternative splicing of the essential exon of murine *Hnrnph1* was reduced by neonatal treatment with 30 µg of *Hnrnph2* ASO in whole brains of juvenile WT and *Hnrnph2* P209L mice. (**F**) Western blot analysis of HEK293T cells transfected with either GFP or GFP-HNRNPH2. Cell lysates were immunoblotted with antibodies against HNRNPH1 (top) and HNRNPH2 (bottom). Note that the HNRNPH1 antibody also recognizes GFP-HNRNPH2, whereas HNRNPH2 antibody only detects overexpressed GFP-HNRNPH2. Endogenous HNRNPH1 protein levels are shown in the accompanying bar graph. (**G**) Schemes for human *HNRNPH2* ASO treatment (10 µg) in human iPSC-derived neurons. (**H**) ddRT-PCR showing levels of *HNRNPH2* and *HNRNPH1* mRNA levels in the three ASO treatment schemes as shown in (G). (**I**) Human iPSC-derived neurons were treated with vehicle, 10 µg of control ASO or human *HNRNPH2* ASO using scheme 3 in G. ddRT-PCR showing levels of *HNRNPH2* and *HNRNPH1* mRNA levels. (**J**) Treatment with human 10 µg *HNRNPH2* ASO in iPSC-derived neurons results in significant increase in HNRNPH1 protein level. Actin was used as a loading control. ns: Non-specific band. \*\*\*\**P* < 0.0001, \*\*\**P* < 0.001, \*\**P* < 0.01, \**P* < 0.05 by 2-way ANOVA with Sidak’s multiple comparison (D and E), student’s t-test (F and J), one-way ANOVA with Dunnett’s multiple comparison (H and I), *n* = 3-4 mice per group (D and E), *n* = 3 technical repeats (F, H-J). Data shown as mean ± SD.

**Table S1.**
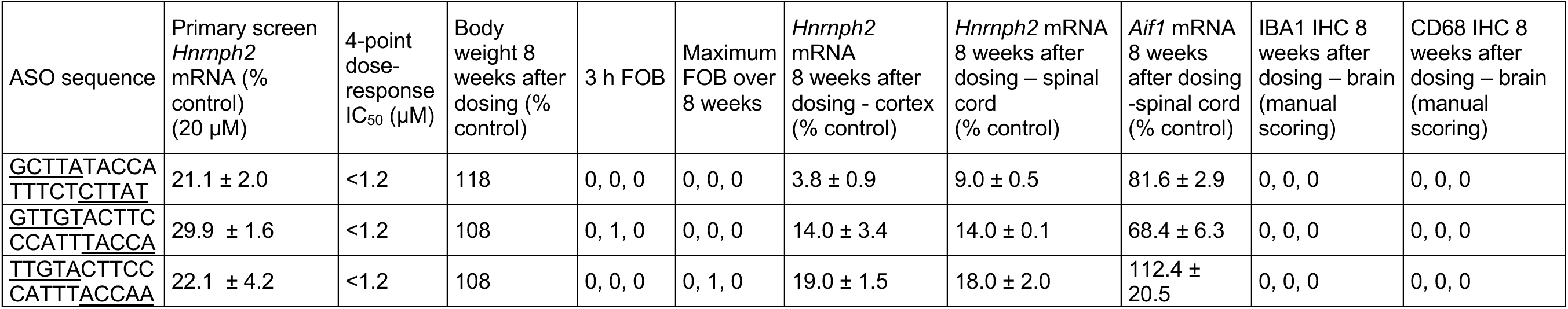
Screening summary of 3 lead mouse *Hnrnph2* ASOs. An example of in vitro and in vivo screening data for 3 lead ASOs designed to target mouse *Hnrnph2*. The ASOs were designed as chimeric 2′-MOE/DNA oligonucleotides with mixed phosphorothioate/phosphodiester linkages. The 2′-MOE nucleotides are underlined. ASOs were initially screened in mouse cortical neurons at 20 µM concentration. The most active ASOs were evaluated in a 4-point dose-response curve ranging in concentration from 1.2 to 32 µM. The most active ASOs were evaluated in female C57/BL6 mice (n = 3) at a dose of 700 µg by ICV injection using a functional observational battery (FOB, scored on a scale of 0 to 7) as described (*3, 4*). Phosphate-buffered saline was used as vehicle control. Eight weeks after injection, mice were sacrificed (n = 3), spinal cord and cortical tissue were isolated, and RNA was extracted. The relative levels of *Hnrnph2* and *Aif1* mRNA were analyzed using qRT-PCR. Brains were also processed for immunohistochemistry (IHC) for IAB1 and CD68 and scored manually for the presence of neuroinflammation (0 = no different from control).

| ASO sequence | Primary screen *Hnrmph2* mRNA (% control) (20 μM) | 4-point dose-response IC<sub>50</sub> (μM) | Body weight 8 weeks after dosing (% control) | 3 h FOB | Maximum FOB over 8 weeks | *Hnrmph2* mRNA 8 weeks after dosing - cortex (% control) | *Hnrmph2* mRNA 8 weeks after dosing – spinal cord (% control) | *Aif1* mRNA 8 weeks after dosing -spinal cord (% control) | IBA1 IHC 8 weeks after dosing – brain (manual scoring) | CD68 IHC 8 weeks after dosing – brain (manual scoring) |
| --- | --- | --- | --- | --- | --- | --- | --- | --- | --- | --- |
| <u>G</u>CTTATACCA<br>TTTCT<u>C</u>TTAT | 21.1 ± 2.0 | <1.2 | 118 | 0, 0, 0 | 0, 0, 0 | 3.8 ± 0.9 | 9.0 ± 0.5 | 81.6 ± 2.9 | 0, 0, 0 | 0, 0, 0 |
| <u>G</u>TTGTACTTC<br>CCATT<u>T</u>ACCA | 29.9 ± 1.6 | <1.2 | 108 | 0, 1, 0 | 0, 0, 0 | 14.0 ± 3.4 | 14.0 ± 0.1 | 68.4 ± 6.3 | 0, 0, 0 | 0, 0, 0 |
| <u>T</u>TGTA<u>C</u>TTCC<br>CATT<u>T</u>ACCAA | 22.1 ± 4.2 | <1.2 | 108 | 0, 0, 0 | 0, 1, 0 | 19.0 ± 1.5 | 18.0 ± 2.0 | 112.4 ± 20.5 | 0, 0, 0 | 0, 0, 0 |

**Table S2.** Pathology (H&E, IHC) of brain regions. C57BL/6J pups were injected via ICV with a single bolus of 30 µg control ASO, *Hnrnph2* ASO, or vehicle at PND 2-4. At 3 weeks of age, whole brains were harvested and evaluated by histology for signs of neuroinflammation or systemic toxicity. Tissues were evaluated for severity with a scale of 0-5. A score of 0 indicates a negative finding (the lesion/feature (H&E) or positive cells (IHC) are absent in the examined slide). A score of 1 indicates a minimal finding (less than 10% based on a visually estimated approximation). A score of 2 indicates a mild finding (11 to 25% based on a visually estimated approximation). Increasing scores are 3 (moderate), 4 (marked), and 5 (severe) (*10*). There were no histopathologic (light microscopic) indicators suggestive of systemic toxicity associated with the administration of 30 µg of control or *Hnrnph2* ASO in any of the organs in any of the six animals. There were no noteworthy neuropathologic findings in the brain, spinal cord, dorsal root ganglia, or spinal nerve roots that could be considered related to the administration of 30 µg of control or *Hnrnph2* ASO in any of the six animals. A mild unilateral needle tract lesion was evident in the cerebral cortex in one animal that was administered 30 µg of control ASO. Focal aggregates of GFAP-positive reactive astrocytes, AIF1/IBA1-positive activated microglia, and CD68-positive macrophages were evident within and around this lesion. Another animal that was administered 30 µg of control ASO had a small, focally extensive, unilateral aggregate of GFAP-positive reactive astrocytes, AIF1/IBA1-positive activated microglia, and CD68-positive macrophages suggestive of a needle tract lesion. However, an overt needle tract lesion was not evident in the brain of this animal. The minimal increase in GFAP-positive astrocytes noted in the adjacent brain regions in these two animals was considered related to overt/suspected needle tract lesions. Needle tract lesions were not evident in the brains of the other four animals. Apart from the lesions noted above, there were no overt neuroinflammatory lesions in any of the brain sections from any of the six animals. However, immunohistochemistry examination revealed minimal to mild increase in AIF1/IBA1-positive activated microglia, and CD68-positive macrophages in the brain in all six animals. Similarly there was minimal to mild increase in the presence of CD68-positive and CD163-positive macrophages in the leptomeninges, choroid plexus, and neuroparenchymal perivascular spaces. These findings were considered related to class-specific, sequence-independent accumulation of macrophages consequent to ASO administration. However, related class-specific effects that are often reported in some nonclinical safety studies such as intra-lysosomal accumulation of ASO cargo (often independent of sequence specificity) seen light microscopically as the accumulation of basophilic granules in renal proximal convoluted tubules and the presence of characteristic vacuolated macrophages in multiple tissues were not evident in any of the six animals.

| Tissue & microscopic feature | 30 µg control ASO |  |  | 30 µg *Hnrnp2* ASO |  |  |
| --- | --- | --- | --- | --- | --- | --- |
|  | Animal 1 | Animal 2 | Animal 3 | Animal 1 | Animal 2 | Animal 3 |
| **Brain (H&E)** Needle tract lesion, cerebral cortex | 0<br>(not present in multiple sections, 3 steps/levels; small focus of hemosiderophages, unilateral, subcallosal, parietal region) | 2<br>(mild, regionally extensive, unilateral, cerebral cortex; additional aggregates of hemosiderophages, leptomeninges, mesencephalic tectum) | 0<br>(not present in multiple sections, 3 steps/levels) | 0<br>(not present in multiple sections, 3 steps/levels) | 0<br>(not present in multiple sections, 3 steps/levels) | 0<br>(not present in multiple sections, 3 steps/levels) |
| **Brain (H&E)** Inflammation, mixed immune cell, neuroparenchyma | 0 | 0 | 0 | 0 | 0 | 0 |
| **Brain (H&E)** Inflammation, mononuclear immune cell (lymphocyte), neuroparenchyma | 0 | 0 | 0 | 0 | 0 | 0 |
| **Brain (H&E)** Necrosis, neuroparenchyma | 0 | 0 | 0 | 0 | 0 | 0 |
| **Brain (H&E)** Vacuolation, neuron, cortex, basal nuclei, hippocampus | 0 | 0 | 0 | 0 | 0 | 0 |
| **Brain (H&E, GFAP)** Cellularity, increased, astrocyte, neuroparenchyma (excluding needle tracts) | 1 | 1 | 0 | 0 | 0 | 0 |
| **Brain (H&E, CD68)** Cellularity, increased, macrophage, neuroparenchyma (excluding needle tracts) | 1 | 1 | 1 | 2 | 1 | 2 |
| **Brain (H&E, CD68, CD163)** Cellularity, increased, macrophage, choroid plexus, leptomeninges, perivascular spaces | 2 | 2 | 2 | 2 | 2 | 2 |
| **Brain (H&E, CD68, CD163)** Aggregation, vacuolated macrophage, choroid plexus, leptomeninges, perivascular spaces | 0 | 0 | 0 | 0 | 0 | 0 |
| **Brain (H&E, IBA1/AIF1)** Cellularity, increased, microglia, neuroparenchyma (excluding needle tracts) | 1 | 2 | 1 | 2 | 1 | 1 |
| **Brain (IHC CD68)** [positive score, neuroparenchyma (excluding needle tracts)] | 1 | 1 | 1 | 2 | 1 | 2 |
| **Brain (IHC CD163)** [positive score, choroid plexus, leptomeninges, perivascular spaces] | 1 | 1 | 1 | 1 | 1 | 1 |
| **Brain (IHC GFAP)** [positive score of GFAP+ cells beyond the extent that would be seen in normal young uninflamed mouse brains] | 1 | 1 | 0 | 0 | 0 | 0 |
| **Brain (IHC IBA1/AIF1)** [positive score of IBA1/AIF1+ cells beyond the extent that | 1 | 2 | 1 | 2 | 1 | 1 |
| would be seen in normal young uninflamed mouse brains] |  |  |  |  |  |  |

**Table S3.** Pathology (H&E, IHC) of all organs. C57BL/6J pups were injected via ICV with a single bolus of 30 µg control ASO, *Hnrnph2* ASO, or vehicle at PND 2-4. At 3 weeks of age, mice were subjected to a standard full necropsy and tissues evaluated by histology for signs of neuroinflammation or systemic toxicity. ASENP = All structures of the middle and inner ear are not available for evaluation in every section/every animal; limited tissue available for examination; ITAE = Insufficient tissue available for evaluation; ITAEC = Insufficient tissue available for evaluation in the second/contralateral eye; NNATMF = No noteworthy ASO-related toxicologically relevant microscopic findings; NPOS/NAE = Not present on the slide/not available for examination; OPAE = one of a pair available for evaluation.

| Tissues | 30 µg control ASO |  |  | 30 µg *Hnrnp2* ASO |  |  |
| --- | --- | --- | --- | --- | --- | --- |
|  | Animal 1 | Animal 2 | Animal 3 | Animal 1 | Animal 2 | Animal 3 |
| Dorsal root ganglia | NNATMF | NNATMF | NNATMF | NNATMF | NNATMF | NNATMF |
| Spinal cord, cervical | NNATMF | NNATMF | NNATMF | NNATMF | NNATMF | NNATMF |
| Spinal cord, lumbar | NNATMF | NNATMF | NNATMF | NNATMF | NNATMF | NNATMF |
| Spinal cord, thoracic | NNATMF | NNATMF | NNATMF | NNATMF | NNATMF | NNATMF |
| Spinal nerve roots | NNATMF | NNATMF | NNATMF | NNATMF | NNATMF | NNATMF |
| Heart | NNATMF | NNATMF | NNATMF | NNATMF | NNATMF | NNATMF |
| Muscle, skeletal, appendicular | NNATMF | NNATMF | NNATMF | NNATMF | NNATMF | NNATMF |
| Trachea | NNATMF | NNATMF | NNATMF | NNATMF | NNATMF | NNATMF |
| Lung | NNATMF | NNATMF | NNATMF | NNATMF | NNATMF | NNATMF |
| Gland, thyroid | NNATMF, OP AE | NNATMF, OP AE | NNATMF, OP AE | NNATMF | NNATMF | NNATMF, OP AE |
| Gland, parathyroid | NPOS/NAE | NPOS/NAE | NPOS/NAE | NPOS/NAE | NPOS/NAE | NNATMF, OP AE |
| Esophagus | NNATMF | NNATMF | NNATMF | NNATMF | NNATMF | NNATMF |
| Kidney | NNATMF | NNATMF | NNATMF | NNATMF | NNATMF | NNATMF |
| Liver | NNATMF | NNATMF | NNATMF | NNATMF | NNATMF | NNATMF |
| Gall bladder | NPOS/NAE | NNATMF | NNATMF | NNATMF | NNATMF | NNATMF |
| Gland, adrenal | NNATMF | NNATMF | NNATMF | NNATMF | NNATMF | NNATMF |
| Thymus | NNATMF | NNATMF | NNATMF | NNATMF | NNATMF | NNATMF |
| Spleen | NNATMF | NNATMF | NNATMF | NNATMF | NNATMF | NNATMF |
| Lymph nodes | NNATMF | NNATMF | NNATMF | NNATMF | NNATMF | NNATMF |
| Gland, mammary | NNATMF | NPOS/NAE | NPOS/NAE | NNATMF | NNATMF | NPOS/NAE |
| Glands, salivary (parotid, mandibular) | NNATMF | NNATMF | NNATMF | NNATMF | NNATMF | NNATMF |
| Stomach | NNATMF | NNATMF | NNATMF | NNATMF | NNATMF | NNATMF |
| Pancreas | NNATMF | NNATMF | NNATMF | NNATMF | NNATMF | NNATMF |
| Small intestines (duodenum, jejunum, ileum) | NNATMF | NNATMF | NNATMF | NNATMF | NNATMF | NNATMF |
| Gut associated lymphoid tissue (GALT) | NNATMF | NNATMF | NNATMF | NNATMF | NNATMF | NNATMF |
| Large intestines (cecum, colon, rectum) | NNATMF | NNATMF | NNATMF | NNATMF | NNATMF | NNATMF |
| Reproductive tract, ovary | NNATMF, OP AE | NNATMF | [male animal] | NNATMF | NNATMF | NNATMF |
| Reproductive tract, oviduct | NNATMF, OP AE | NNATMF | [male animal] | NNATMF | NNATMF | NNATMF |
| Reproductive tract, uterus/cervix | NNATMF | NNATMF | [male animal] | NNATMF | NNATMF | NNATMF |
| Reproductive tract, vagina | NNATMF | NNATMF | [male animal] | NNATMF | NNATMF | NNATMF |
| Testes | [female animal] | [female animal] | NNATMF immature<br>(no spermatogenesis yet) | [female animal] | [female animal] | [female animal] |
| Epididymis | [female animal] | [female animal] | NNATMF | [female animal] | [female animal] | [female animal] |
| Gland, prostate | [female animal] | [female animal] | NNATMF | [female animal] | [female animal] | [female animal] |
| Seminal vesicles | [female animal] | [female animal] | NPOS/NAE | [female animal] | [female animal] | [female animal] |
| Urinary bladder | NNATMF | NPOS/NAE | NNATMF | NNATMF | NPOS/NAE | NNATMF |
| Skin | NNATMF | NNATMF | NNATMF | NNATMF | NNATMF | NNATMF |
| Eyes | NNATMF | NNATMF, ITAE2 | NNATMF | NNATMF | NNATMF | NNATMF |
| Gland, Harderian | NNATMF | NNATMF | NNATMF | NNATMF | NNATMF | NNATMF |
| Gland, pituitary | NNATMF | NNATMF | NPOS/NAE | NPOS/NAE | NPOS/NAE | NNATMF |
| Head, cranial nerves and ganglia | NNATMF | NNATMF | NNATMF | NNATMF | NNATMF | NNATMF |
| Head, ear | NNATMF | NNATMF | NNATMF | NNATMF | NNATMF | NNATMF |
| Head, nasal cavity | NNATMF | NNATMF | NNATMF | NNATMF | NNATMF | NNATMF |
| Head, oral cavity | NNATMF | NNATMF | NNATMF | NNATMF | NNATMF | NNATMF |
| Head, teeth | NNATMF | NNATMF | NNATMF | NNATMF | NNATMF | NNATMF |
| Head, tongue | NNATMF | NNATMF | NNATMF | NNATMF | NNATMF | NNATMF |
| Nerve, optic | NNATMF | NNATMF | NNATMF | NNATMF | NNATMF | NNATMF |
| Muscle, skeletal, axial | NNATMF | NNATMF | NNATMF | NNATMF | NNATMF | NNATMF |
| Sternum | NNATMF | NNATMF | NNATMF | NNATMF | NNATMF | NNATMF |
| Bone marrow (H&E) | NNATMF | NNATMF | NNATMF | NNATMF | NNATMF | NNATMF |
| Femur | NNATMF | NNATMF | NNATMF | NNATMF | NNATMF | NNATMF |
| Joint, femorotibial | NNATMF | NNATMF | NNATMF | NNATMF | NNATMF | NNATMF |

**Table S4.**
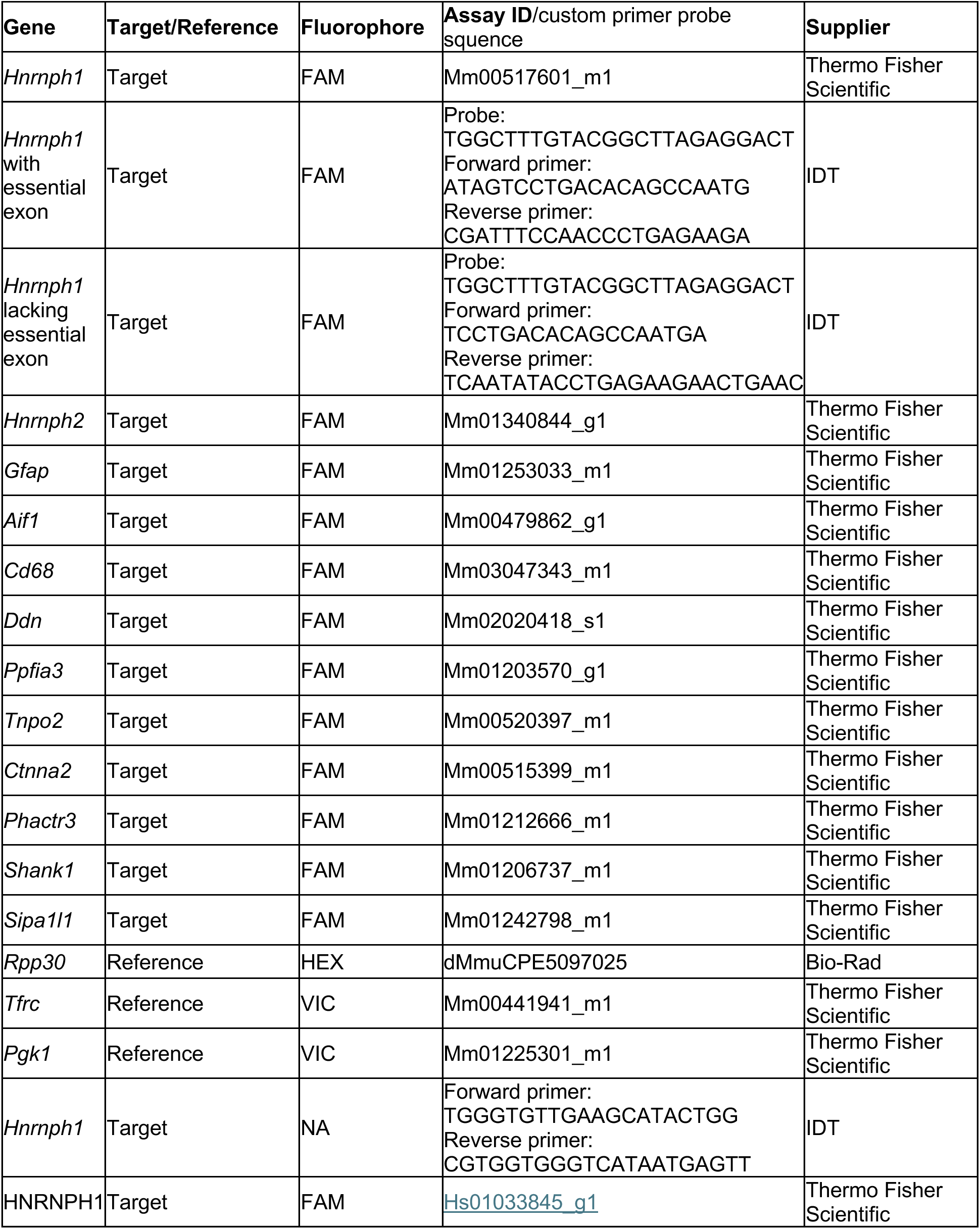
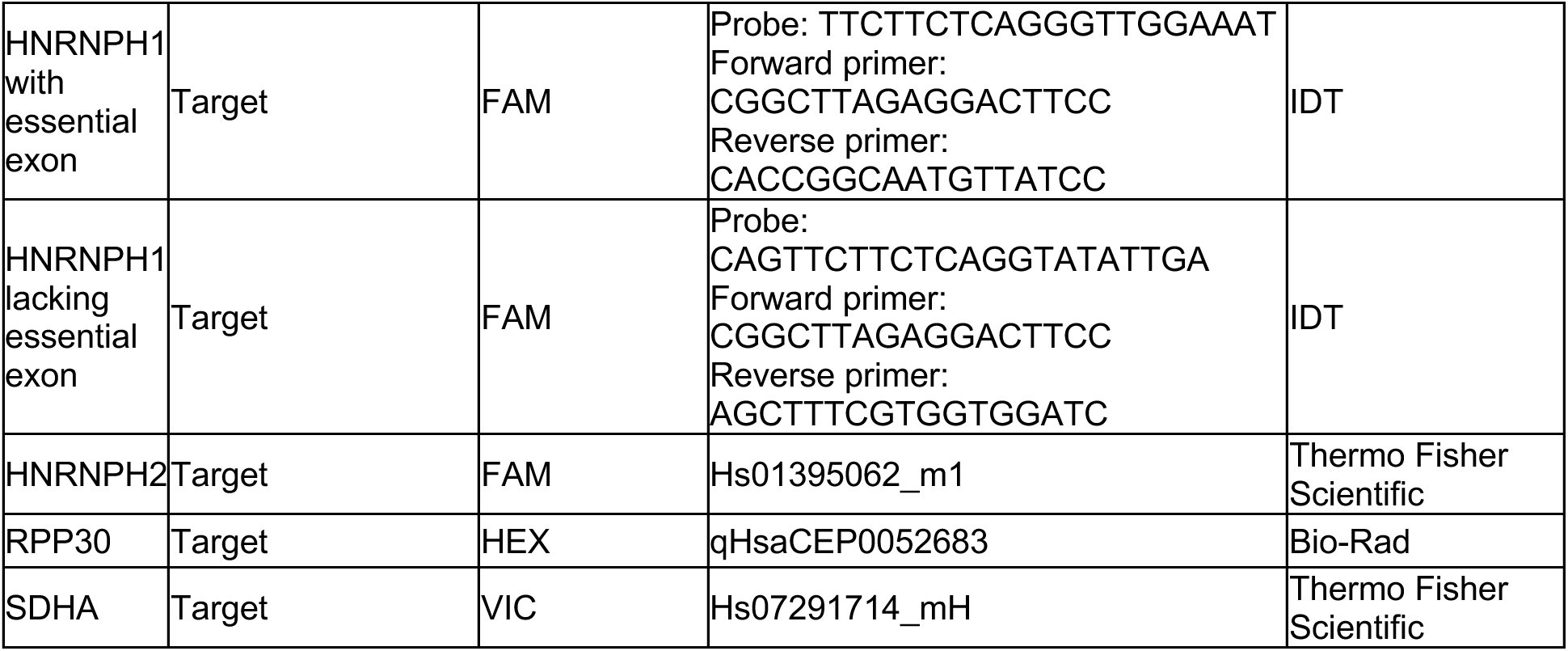
Gene expression assays used for mouse brain ddRT-PCR and RT-PCR, and human ddRT-PCR.

**Table S5.** Antibodies used in western blots.

| Antibody | Specificity | Catalog number | Supplier | Host | Dilution used |
| --- | --- | --- | --- | --- | --- |
| Anti-HNRPH2 antibody [EPR12170(B)] | Specific for HNRNPH2 (mouse only, detects both human HNRNPH2 and HNRNPH1) | ab179439 | Abcam | Rabbit | 1:500 |
| hnRNP H1 polyclonal Antibody | Specific for HNRNPH1 (mouse and human) | PA5-50678 | Thermo Fisher Scientific | Rabbit | 1:1000 |
| GAPDH (D4C6R) Mouse mAb | GAPDH (cytoplasmic protein) | #97166 | Cell Signaling Technology | Mouse | 1:2000 |
| HDAC2 Monoclonal Antibody (HDAC2-62) | HDAC2 (nuclear protein) | MA1-25451 | Thermo Fisher Scientific | Mouse | 1:1000 |
| β-Actin (13E5) Rabbit mAb | Total β-actin | 4970 | Cell Signaling Technology | Rabbit | 1:1000 |
| HA tag antibody | HA tag | NB600-363 | Novus Bio | Rabbit | 1:1000 |
| Anti-GFP | GFP tag | 11814460001 | Millipore Sigma | Mouse | 1:1000 |
| Secondary ab's for human WBs |  |  |  |  |  |
| IRDye 800CW Donkey anti-Rabbit IgG Secondary Antibody | Heavy & light chains of rabbit IgG, and light chains common to most rabbit immunoglobulins | 926-32213 | LICOR | Donkey | 1:10,000 |
| IRDye 680RD Goat anti-Mouse IgG Secondary Antibody | Heavy & light chains of mouse IgG<sub>1</sub>, IgG<sub>2a</sub>, IgG<sub>2b</sub>, and IgG<sub>3</sub>, and light chains of mouse IgM and IgA | 926-68070 | LICOR | Goat | 1:10,000 |

**Table S6.**
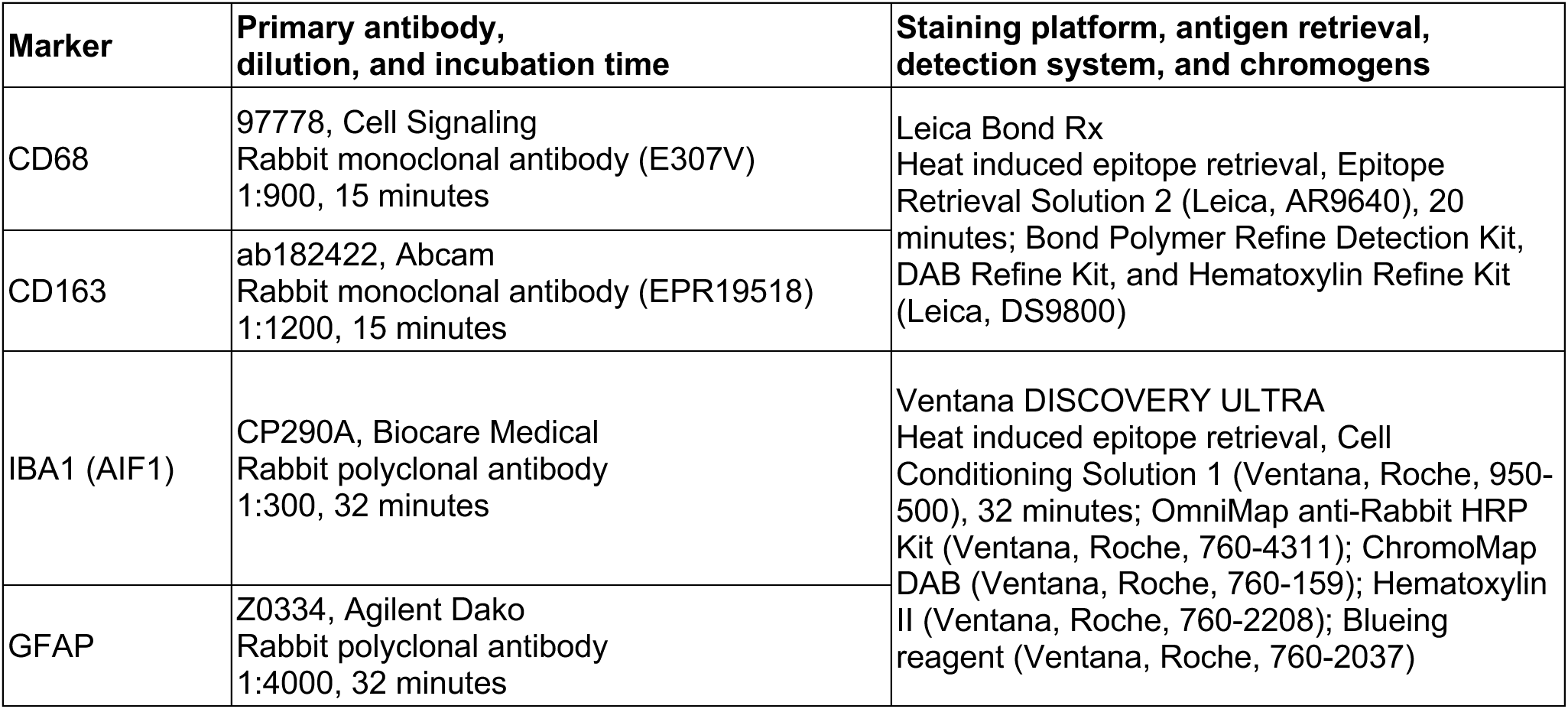
Epitope retrieval, primary antibodies, staining platform, and detection system used in histopathology.

| Marker | Primary antibody, dilution, and incubation time | Staining platform, antigen retrieval, detection system, and chromogens |
| --- | --- | --- |
| CD68 | 97778, Cell Signaling<br>Rabbit monoclonal antibody (E307V)<br>1:900, 15 minutes | Leica Bond Rx<br>Heat induced epitope retrieval, Epitope Retrieval Solution 2 (Leica, AR9640), 20 minutes; Bond Polymer Refine Detection Kit, DAB Refine Kit, and Hematoxylin Refine Kit (Leica, DS9800) |
| CD163 | ab182422, Abcam<br>Rabbit monoclonal antibody (EPR19518)<br>1:1200, 15 minutes |  |
| IBA1 (AIF1) | CP290A, Biocare Medical<br>Rabbit polyclonal antibody<br>1:300, 32 minutes | Ventana DISCOVERY ULTRA<br>Heat induced epitope retrieval, Cell Conditioning Solution 1 (Ventana, Roche, 950-500), 32 minutes; OmniMap anti-Rabbit HRP Kit (Ventana, Roche, 760-4311); ChromoMap DAB (Ventana, Roche, 760-159); Hematoxylin II (Ventana, Roche, 760-2208); Blueing reagent (Ventana, Roche, 760-2037) |
| GFAP | Z0334, Agilent Dako<br>Rabbit polyclonal antibody<br>1:4000, 32 minutes |  |

**Table S7.** Antibodies used for mouse brain immunofluorescence.

| Antibody | Host | Specificity | Dilution used | Catalog number | Supplier |
| --- | --- | --- | --- | --- | --- |
| Anti-ASO antibody | Rabbit | ASO phosphorothioate backbone | 1:20,000 | NA | Ionis |
| Anti-HNRPH2/HNRNPH2 antibody [EPR12171] | Rabbit | Specific for HNRNPH2 | 1:100 | ab181171 | Abcam |
| Anti-GFAP antibody | Rabbit | GFAP (astrocyte marker) | 1:1000 | Z033429 | Agilent |
| Anti-GFAP monoclonal antibody (2.2B10) | Rat | GFAP (astrocyte marker) | 1:1000 | 13-0300 | Thermo Fisher Scientific |
| Anti-NeuN antibody | Guinea pig | NeuN (neuronal marker) | 1:250 | 266004 | Synaptic Systems |
| Anti-IBA1 antibody | Guinea pig | IBA1 (microglial marker) | 1:500-1000 | 234308 | Synaptic Systems |
| Anti-Carbonic Anhydrase II antibody | Sheep | Ca2 (oligodendrocyte marker) | 1:500 | AHP206 | Bio-Rad |
| Anti-Calbindin D-28K antibody | Chicken | Calbindin (Purkinje cell marker) | 1:1000 | NBP2-50028 | Novus Biologicals |
| Anti-Parvalbumin antibody | Guinea pig | Parvalbumin (Purkinje cell / interneuron marker) | 1:500 | GP72 | Swant |
| Alexa Fluor 488 anti-rabbit antibody | Donkey | IgG | 1:500 | 711-547-003 | Jackson ImmunoResearch |
| Alexa Fluor 488 anti-rabbit antibody | Donkey | IgG (Fab) | 1:500 | 711-545-152 | Jackson ImmunoResearch |
| Alexa Fluor 568 anti-guinea pig antibody | Donkey | IgG | 1:500 | A11075 | Thermo Fisher Scientific |
| Alexa Fluor 568 anti-rat antibody | Donkey | IgG (Fab) | 1:500 | ab175708 | Abcam |
| Alexa Fluor 594 anti-sheep antibody | Donkey | IgG | 1:500 | 713-585-003 | Jackson ImmunoResearch |
| Alexa Fluor 647 anti-chicken antibody | Donkey | IgY | 1:500 | 703-605-155 | Jackson ImmunoResearch |
| Alexa Fluor 647 anti-guinea pig antibody | Donkey | IgG | 1:500 | 706-605-148 | Jackson ImmunoResearch |
| Alexa Fluor 647 anti-rat antibody | Donkey | IgG | 1:500 | 712-605-153 | Jackson ImmunoResearch |
| Alexa Fluor 750 anti-guinea pig antibody | Goat | IgG | 1:500 | ab175745 | Abcam |

